# Implication of structural constrains facilitating the functional evolution of *Pseudomonas aeruginosa* KPR2 into a versatile α-keto acid reductase

**DOI:** 10.1101/2024.01.05.573888

**Authors:** Gourab Basu Choudhury, Saumen Datta

## Abstract

Protein structure and function dynamics in molecular evolution are intertwined. Theoretical concepts linking structure, function, and evolution of a protein, while often intuitive, necessitate validation through investigations in real-world systems. Our study empirically explores the implications of multiple panE2 gene copies in an organism, shedding light on the functional roles and evolutionary trajectories of *Pseudomonas aeruginosa’s* second copy of Ketopantoate reductase (PaKPR2) and its inactivity against the natural substrate Ketopantoate. Evolutionary changes in functional traits were examined around the active site through crystal structures and biochemical analysis. Primarily, apoKPR2 structures reveal a transformed active site cleft, forming a two-sided pocket, while substrate entry is regulated by a molecular gate. Despite cleft closure, molecular interaction properties and activity analysis of PaKPR2 suggest that it can be a versatile keto-acid reductase. However, detailed structural insights from the ligand-bound binary complex of PaKPR2-NADPH and PaKPR2-Ketoisoleucine reveal that the ligand-binding interactions at the active site are conserved and restricted to the molecules of appropriate shape and size that can be accommodated in the available space. Finally, a ternary complex structure, PaKPR2-NADP+-KIC, was solved to understand its functional evolution in terms of the residue microenvironment at the catalytic site. Collectively, the results give detailed visual experiences of different structural perspectives of the protein’s functional evolution.

## Introduction

*Pseudomonas aeruginosa* is an opportunistic pathogen ubiquitous in nature, which is shown in its adaptable metabolic capacity and wide range of attributes for environmental adaptability. As a significant causative agent of Cystic fibrosis, *P. aeruginosa* has been found to opt for altered phenotypes supported with different metabolic requirements in case of changing its developmental niche from the natural environment to the host system (Palmer et al, 2007, Turner et al, 2015). The organism has a large genome size compared to other prokaryotes and displays high genomic diversity and complexity, consistent with its ubiquitous environmental adaptability and resistance to multiple drugs (Stover, et al, 2000, Weiser et al, 2019). Evolution is driven by natural selection in complex and dynamic environmental conditions, leading to intricate and sometimes unconventional directions that impact numerous genes and metabolic solutions (La Rosa et al, 2018).

Long-studied genome evolution and adaptation of bacterial populations provided evidence related to the importance of their Genomic diversity and dynamics (Gray and Fitch, 1983, Yoshikuni et al., 2006). Dynamics implicates the fundamental processes by which microbes gain or lose functions that are deeply responsible for their evolution (Wiedenbeck and Cohan, 2011; Blount et al., 2012). They also contribute to the adaptability of microorganisms to various environmental niches (Bratlie et al., 2010; Hottes et al., 2013). Through gene duplication and divergence, a significant number of enzymes have evolved, while horizontal gene transfer and further dynamics have resulted in the differential fates of the bacterial genes. (Serres et al., 2009; Wang and Chen, 2018). Prokaryotic genome evolution is drastically contributed by Horizontal Gene Transfer (HGT) (Kunin et al, 2003). Comparative genomics with flux balance analysis has shown that in the past 100 million years most changes to the metabolic network of Escherichia coli are due to gene duplication and horizontal gene transfer. Interestingly, the role of the latter is major in metabolic pathway evolution (Koonin et al., 2002). The accusation of genes via HGT may result in producing xenologous genes, which are, in the true sense, homologs to other genes present in the same genome. Eventually, these xenologoues under selection stress may gain new functions (neofunctionalization) or lose their existing function in a broad evolutionary window (Bridgham et al, 2008).

One of the well-studied metabolic pathways is pantothenate synthesis, concerning genomic, structural and functional context. Our recent research revealed that the genes responsible for the vitamin B5 production pathway are subject to dynamic evolutionary selection, resulting in the acquisition of multiple copies of these genes by several bacterial species or their dependence on alternative sources of this crucial nutrient. KPR is one of the genes involved in the production of pantothenate that is reported to be duplicated in around 20% of bacterial species. panE gene codes for the enzyme 2-dehydropantoate reductase or Ketopantoate reductase (KPR) that takes part in the second step of the pantothenate synthesis pathway, where it converts ketopantoate to pantoate. Interestingly in *the P. aeruginosa* genome, there are two reported genes for this conversion step according to the KEGG database with different Gene IDs having sequence similarity of 34% and sequence Identity of 24% between them. However, their role and functional characteristics are yet to be discovered fully. Functional and Structural perspective studies of the panE1 established its efficiency as canonical Ketopantoate reductase in the organism, however, nothing much is known about the second copy of the KPR i.e. PanE2. These KPRs in *P. aeruginosa* did not cluster together, so, it is evident that the duplicate copies of KPRs do not follow the traditional 16 s rRNA-based phylogenetic tree. From previously reported phylogenetic analysis it is understood that panE1 and *Azotobacter chroococcum* KPR have a greater nucleotide sequence similarity than does *P. aeruginosa* KPR2 (PaKPR2). That presumed that the panE1 gene is most likely acquired horizontally since it lacks a tight evolutionary link and has a greater sequence similarity with the panE genes of *A. chroococcum.* In contrast to *P.aeruginosa*, a higher similarity was shown between different copies of KPRs from *E. faecalis*, and they also coexisted in the same phylogenetic tree. Therefore, it appears that both *P. aeruginosa* and *A. chroococcum* obtained numerous copies of KPRs by horizontal gene transfer events whereas *E. faecalis* most likely got multiple copies of KPRs through genomic duplication events (Khanppnavar et al,2019).

All these genome dynamics lead to the more evolved and diverse genomic context in the organism that allows them to sustain in different external conditions depending on their life cycles (Hooper, 2003, Klein et al., 2009, Khademi et al., 2019). Apart from the regulation and altered function, the divergence of the gene also leads to differential subcellular localization, change of function, and evolution of catalytic mechanism (Copley et al, 2020). Hence the multiple copies of isoenzymes with the same function in an organism indicate something more along with their native state of function. Analysing their present functional state in the context of structural notion can elaborate vast fields of uncertainty related to their advantageous or depreciating effects on the population (Rocha, 2006, Lovell et al., 2010).

KPR is a member of the 6-phosphogluconate dehydrogenase superfamily, which is a subset of the incredibly varied and extensive group of Rossmann-like NAD(P)-binding dehydrogenases. Other enzymes like acetohydroxy acid isomeroreductase and short-chain L-3-hydroxyacyl-CoA dehydrogenase are members of the 6-phosphogluconate dehydrogenase superfamily. Therefore, it is necessary to discriminate between true and other enzymes. Recently, it has been discovered that Enterococcus faecalis IAM10071 and Lactococcus lactic NADH-dependent D-2-hydroxy acid dehydrogenases are mistakenly classified as PanE (KPR) (Wada et al., 2008; Chambellon et al., 2009).

Regarding the connections between structure and evolution, we have a variety of theoretical concepts. Even if many of these are quite intuitive and have a “sense” of relevance, real relevance must be established by investigating systems in the real world (Rocha and Danchin, 2004, Schlicker, et al., 2007, Franzosa et al., 2008). Keeping this in mind, our study has illustrated existing perspectives of the evolution by showing structural details at the atomic level with different functional states. of a metabolic enzymes PanE2 of *P. aeruginosa* that lack activity against Ketopantoate but gained broad substrate specificity against several other alpha keto acids like Pyruvate and glyoxylate, Ketoisocaproate,3methyl 2 oxo ketopentanoate, 4-phenyl 2 oxo butanoate, phenyl pyruvate, 2keto-hexanoate etc. Additionally, the enzyme kinetic assay and ligand interaction analysis employing biophysical methods, elucidated that PanE2 possesses varying affinity for different structurally related alpha-keto acids substrates. Considering these observed phenomena, factors influencing the enhanced functional traits were investigated at the molecular level in terms of residue microenvironment at and around the active site of the protein through several crystal structures of apoKPR2, ligand-bound binary and ternary protein complexes.

Collectively, our study tried to focus on the current status of the second copy of *P. aeruginosa* KPR in the context of their structural divergence and functional diversification which will shed light on the functional role and the metabolic specialization along with other evolutionary constraints of the multiplication event widely spread in different organism for this enzyme.

## 2. RESULTS

### 2.1 Native structures of Ketopantoate reductase 2 of *P. aeruginosa* revealed the possible reason behind the inactivity of the protein towards the Ketopantoate

In many bacterial species, the panE gene, which codes for ketopantoate reductase (KPR) is either missing or present in numerous copies due to dynamic evolutionary selection. To provide insight into the causes of these dynamic evolutionary selects, it would be prudent to examine the conformational dynamics and structural characteristics of multiple copies of KPRs present in same organism as Xeologues. *P. aeruginosa* KPR xenologues has been found to have 34%similarity and 24% identity in their sequences as compared in BLAST analysis (Altschul, S.F., 1990). Ketopantoate reductase enzymes are well characterised in different organism. Their structural significance and functional role in pantothenate pathway are quite elusive from previous extensive studies mainly *in E. coli* (Lobley, C. M, 2005; Ciulli, A., 2007). However, in PDB approx. 23 structures are available from 12 species. But none of these previously highlighted the nature of the xenologues. However, those studies shows that the KPR homologue structures are composed of mainly two globular domains which forms a cleft like junction in between contributing to the formation of the cofactor and substrate binding site. N-terminal domain mainly harbours conserved Rossman fold dedicated for cofactor binding and the C terminal helical domain which in some species found to contribute for dimer formation. Variations in oligomeric state of the enzyme has been reported in different organisms. Such as *Staphylococcus aureus, Ralstonia eutropha, Ralstonia solanacearum, Mycobacterium tuberculosis, Enterococcus faecalis, Methylococcus capsulatus,* and *Bacillus subtilis* were reported to have dimeric KPRs only (PDB entries 4YCA, 3HWR, 3GHY, 4OL9, 2EW2, 3I83, and 3EGO, respectively). But, others like *E. coli, Geobacter metallireducens* and *Porphyromonas gingivalis* and our previously characterised *Pseudomonas aeruginosa KPR1* (PDB entries 1KS9, 3HN2, 2QYT and 5ZIK respectively), was found to be a monomeric unit. Molecular wight difference between the two PaKPR proteins in *Pseudomonas aeruginosa* are very less, one being 34KDa and another 33 KDa. But The elution volume of the Gel filtration profile suggests the PaKPR2 is much bigger in size than the PaKPR1 and compare to standard curve of know mol weight protein it comes between 120 -60 kDa (Supplementary figure 1B and 1C). Moreover, crosslinked protein with 0.025% of glutaraldehyde also analysed to check the molecular weight of the protein in solution. And when the SDS gel band was compared with the untreated sample it shows size variation and the crosslinked product band appeared much above the control around 70kd region of the protein marker which implicates it to be a dimer. (Supplementary figure 1A). It is to be noted here that monomeric or dimeric KPR is already reported in different organisms and they found those proteins functional too. (Ciulli, A., 2007; Sanchez, J. E., 2015; Aikawa, Y., 2016). However, two types of KPR with two different oligomers in specific organism is first time reported here for *P. aeruginosa*.

Interestingly, Dimeric Xenologue KPR of *P.aeruginosa*, which is denoted as PaKPR2/PanE2 in this study, is found to be inactive in an initial activity assay in the presence of NADPH and the substrate Ketopantoate. These observations make a popular choice to identify this gene as a pseudoenzyme with respect to the pantothenate pathway. Because, an enzyme and pseudoenzyme pair typically have relatively little pairwise sequence similarity, despite their extremely high structural similarity. In addition to that, pseudoenzymes share the same superfamily and tertiary structures as the true enzymes, but they lack catalytic residues or have steric disruption at the catalytic site entrance, rendering them catalytically inert (Goldberg, T., & Sreelatha, A. 2023). However, paKPR2 have maintained its active site residues conserved. We have established that from the multiple sequence alignment of PaKPR2 with the sequences of all the previously reported KPR structure. The ketopantoate to pantoate conversion process is primarily facilitated by two lysine residues (E. coli K72 and K176) that are entirely conserved, as well as four conserved acidic residues (E. coli E210, E240, D248, and E256) that could serve as generic acid/base groups. These conserved residues have been highlighted in the MSA results given in supplementary figure 2. To investigate the probable cause behind the inactivity in the context of structural divergence we wanted to find out the structural differences in the specific PaKPR2 protein.

To shed light on the conserved and diversified facets in catalytic mechanism of the protein first thing was to identify the divergences in the native structural features of the protein. We Crystallized it and solved the structures with two different native 3D structures (Figure 1). These structures were deposited in the PDB as 8IWG and 8IWQ (resolution: 2.15 Å and 2.19 Å).It was noted that the secondary structure components like helix and Beta sheets are more or less equal as compared to previously reported KPR homologues in PaKPR2. But the tertiary structure was dimeric, where two C terminal domains of different monomeric unit buried themselves in as end to end joining made the dimeric interface of the protein (highlighted region in Figure 1A). PISA analysis (E. Krissinel and K. Henrick, 2007) of the native structures shows that the interface between the molecules buries 946 Å^2^ of surface area per monomer and includes 25 residues from each chain. The analysis gives Complex Formation Significance Score (CSS) that ranges from 0 to 1 as interface relevance to complex formation which was noted as maximum in this case. Achieved CSS of 0.599 implies that the interface is due to dimer formation. The dimer interface predicts a favourable solvation free energy gain (ΔiG = − 9.9 kcal mol−1) and a P value of 0.125; P values of <0.5 indicate a strong likelihood that the surface is interaction-specific and not because of crystal contacts. Important hydrogen bonds and salt bridges at the dimeric interphase are tabulated in the Table 1 and 2.

**Figure 1:**
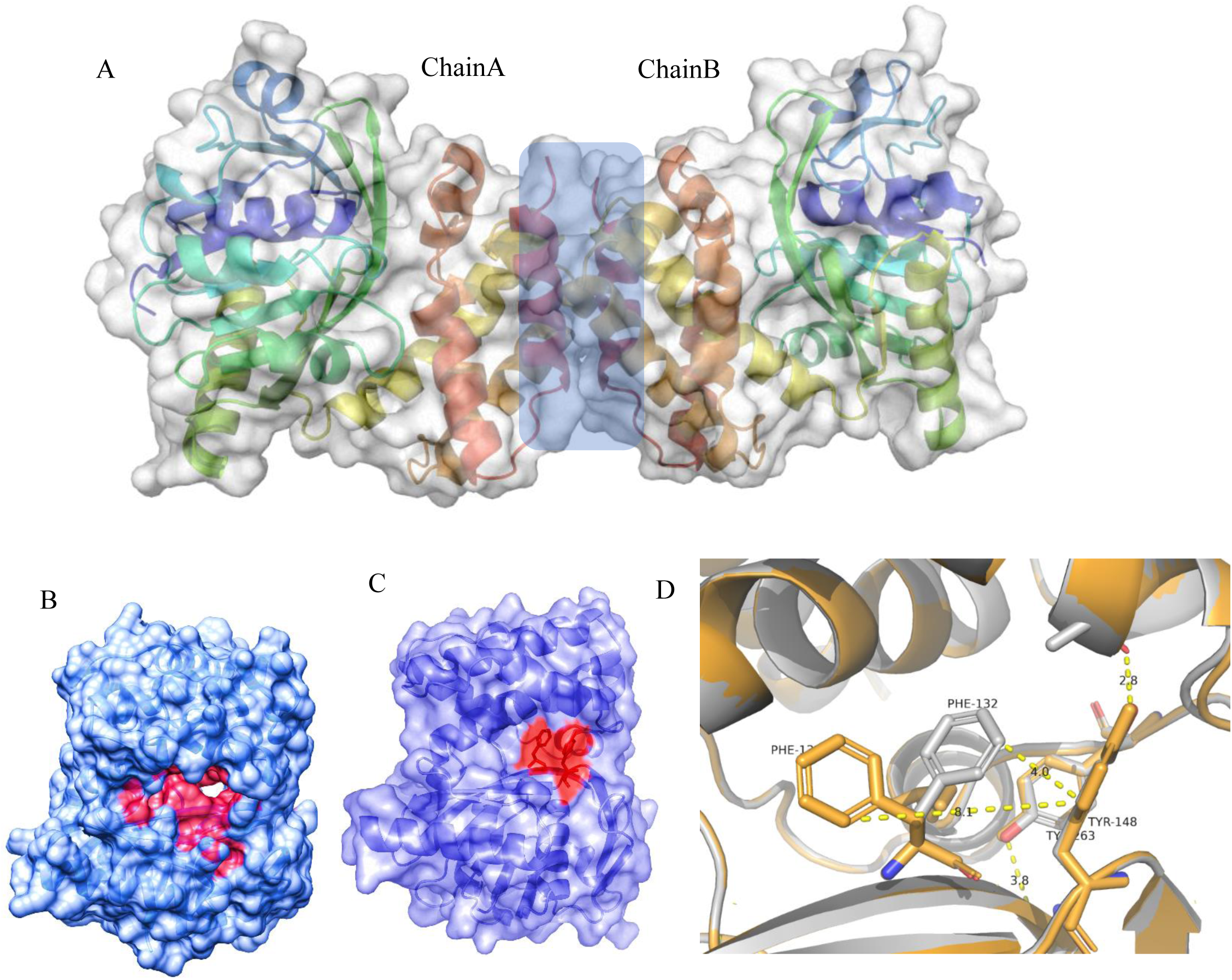
Native structure of PaKPR2. (A)Dimeric KPR with Interface region highlighted with Blue colour box. (B) PDB: 8IWQ Native structure of PaKPR2 with open gate conformation. (C) PDB: 8IWG Native structure of PaKPR2 with closed gate conformation (Gate residues F132 and Y148 showed in red). (D) Two native structure Super imposed evaluate change of orientation of F132 responsible for dynamic molecular gate.

**Table 1:**
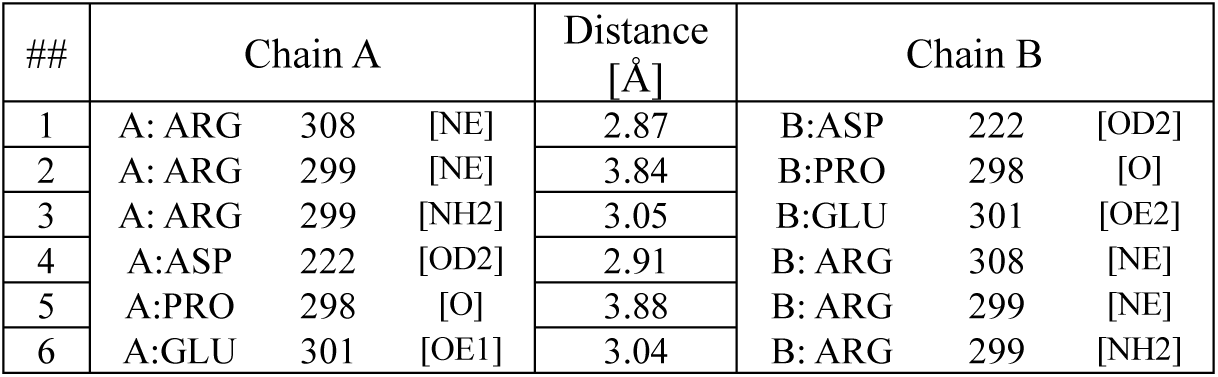
Hydrogen Bonds in Dimer Interface.

**Table 2:** Salt Bridges at Dimer Interface.

| ## | Chain A |  |  | Distance<br>[Å] | Chain B |  |  |
| --- | --- | --- | --- | --- | --- | --- | --- |
| 1 | A: ARG | 308 | [NE] | 3.11 | B:ASP | 222 | [OD1] |
| 2 | A: ARG | 308 | [NE] | 2.87 | B:ASP | 222 | [OD2] |
| 3 | A: ARG | 308 | [NH2] | 3.60 | B:ASP | 222 | [OD2] |
| 4 | A: ARG | 299 | [NE] | 3.89 | B:GLU | 301 | [OE2] |
| 5 | A: ARG | 299 | [NH2] | 3.05 | B:GLU | 301 | [OE2] |
| 6 | A:ASP | 222 | [OD1] | 2.96 | B: ARG | 308 | [NE] |
| 7 | A:ASP | 222 | [OD2] | 2.91 | B: ARG | 308 | [NE] |
| 8 | A:ASP | 222 | [OD2] | 3.43 | B: ARG | 308 | [NH2] |
| 9 | A:GLU | 301 | [OE1] | 3.99 | B: ARG | 299 | [NE] |
| 10 | A:GLU | 301 | [OE1] | 3.04 | B: ARG | 299 | [NH2] |

As we focus more intricate analysis for each monomeric unit, tertiary structure seems a bit squeezed between the two well-constructed domains. It was also observed from our crystal structures that solvent accessibility of the area has been changed to make the active site of the protein more confined. The characteristic hinge region is missing too. Comparative analysis with other reported KPR structure for this hinge region has been shown in the supplementary figure 4. Changes in the residue interaction networks of that area converted the conserved cleft like active side as a two-sided pocket, one for the cofactor entry and another for the substrate entry. Specifically, a HELIX-TURN-HELIX motif of the C-terminal domain comes closer to the opposite β-sheet region present in the N-terminal domain. On a closer look to our apoKPR structures it was found that a tyrosine molecule at position 148 makes an inter domain hydrogen bonds with the carboxyl oxygen of the residue Ala 256 which made the earlier mentioned cleft partially closed (Supplementary Video 1). This domain movement and cleft closue at native form has been found very specific for this protein as compared to other reported KPR native structures as shown by super imposing those structures with PaKPR2 presented in the supplementary figure 5. Molecular evolution has been found to be correlated with certain structural properties, such as solvent exposure, both at the residue and whole-protein levels. Proteins with residues buried in their cores have a higher chance of being preserved throughout evolution than counterparts exposed to solvents (Goldman et al. 1998; Bustamante et al. 2000; Choi et al. 2006; Conant and Stadler 2009). Figure 2 A, 2 B depicts the comparative analysis of the cleft volumes by CastP webserver (Dundas, J, 2006) between two PaKPR clearly shows volume reduction at the space. Even after cofactor binding the space remains higher than the closed counterpart (Figure 2B). In addition to that, in this region one important conformational difference was noted between two native structures. In the 8IWG structure the one end of the cleft the two aromatic residues Phe 132 and Tyr148 mad *π π* interaction and hence no pocket is visible (Figure 1C). However, in the second native structure (8IWQ) where Glycerol was added in higher percentage in cryo-solution during crystallography data collection a 90° rotameric shift of the phenyl group of F132 residue was captured. The outward movement of the sidechain opened the cleft and created a pocket where glycerol made an entry inside it. Hence the structure shows a pocket at the exact place of F132-Y148 interaction site and the region where glycerol has been found is expected to be the ligand binding site (Figure 1C). Relative orientation and corresponding distance of the two residues F132 and Y148 in two different apoKPR structure has been shown in the Figure 1D. We revalidated our claim with the omit maps of the residues in different orientations presented in the supplementary figure 6. It is evident that the space remaining inside the gate after the closure is very less but certainly it was easier to accommodate molecules like glycerol. This gives a clue that the protein might not be a fully dead variant of the Ketopantoate reductase in the organism. If we consider the molecular structure of Ketopantoate (Mol wt. 161.133), it has two-legged methyl group at C3 that makes it a little bulky at the middle (Supplementary figure 3) that cause the active site residues of the PaKPR2 to employ steric hindrances to accommodate the molecule inside it. Else for the bigger size its entry is restricted by the molecular gate created by Y148 and F 132 of the protein. It seems that the cleft has been converted into a two-way pocket or tunnel, having a substrate entry site and another cofactor entry site.

**Figure 2:**
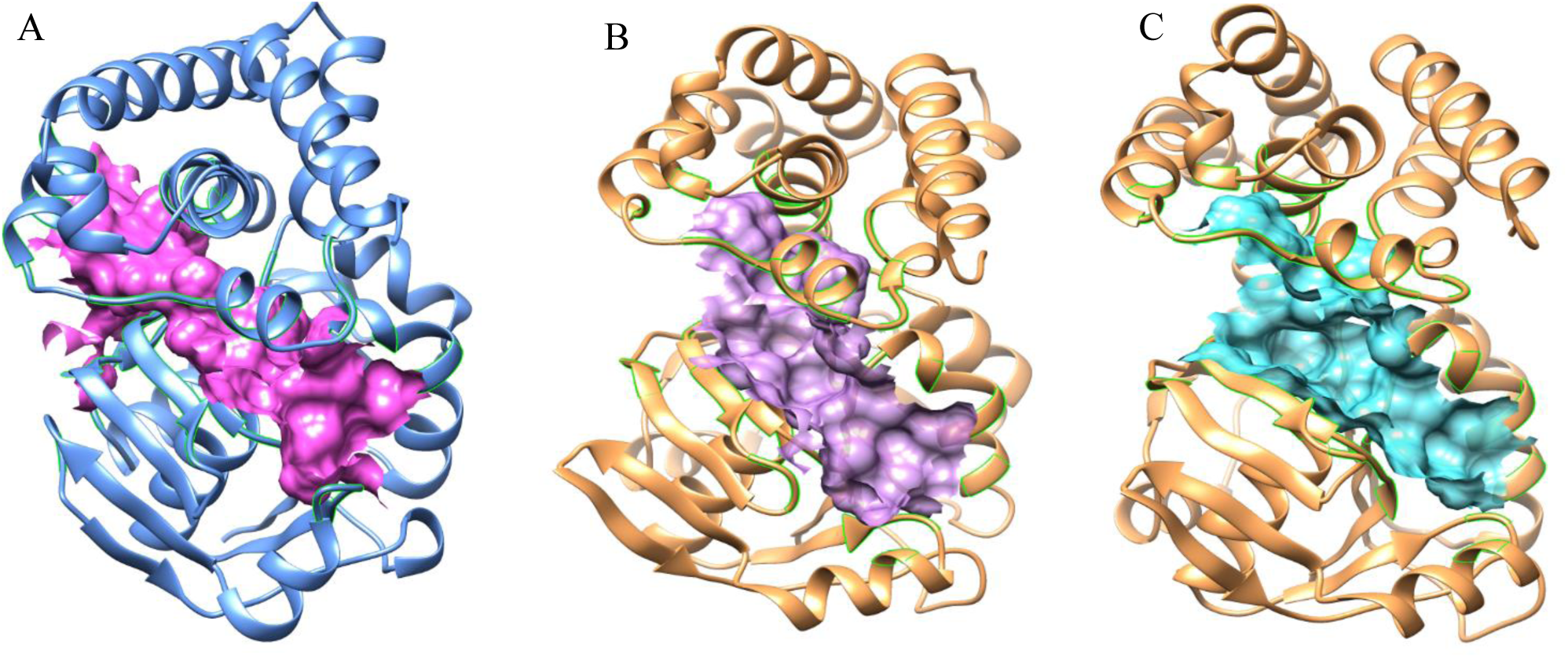
Surface representation of partially closed Cleft turned Pocket volume of (A) KPR2 (8IWG) (volume 2756 Å_3_ ) compared to open cleft of (B) apo KPR1 (5ZIK) ( volume 3144 Å_3_ ) and (c) NADPH-KPR1 (5ZIX) ( volume 2851 Å_3_ ) structure of *Pseudomonas aeruginosa*.

**Figure 3.**
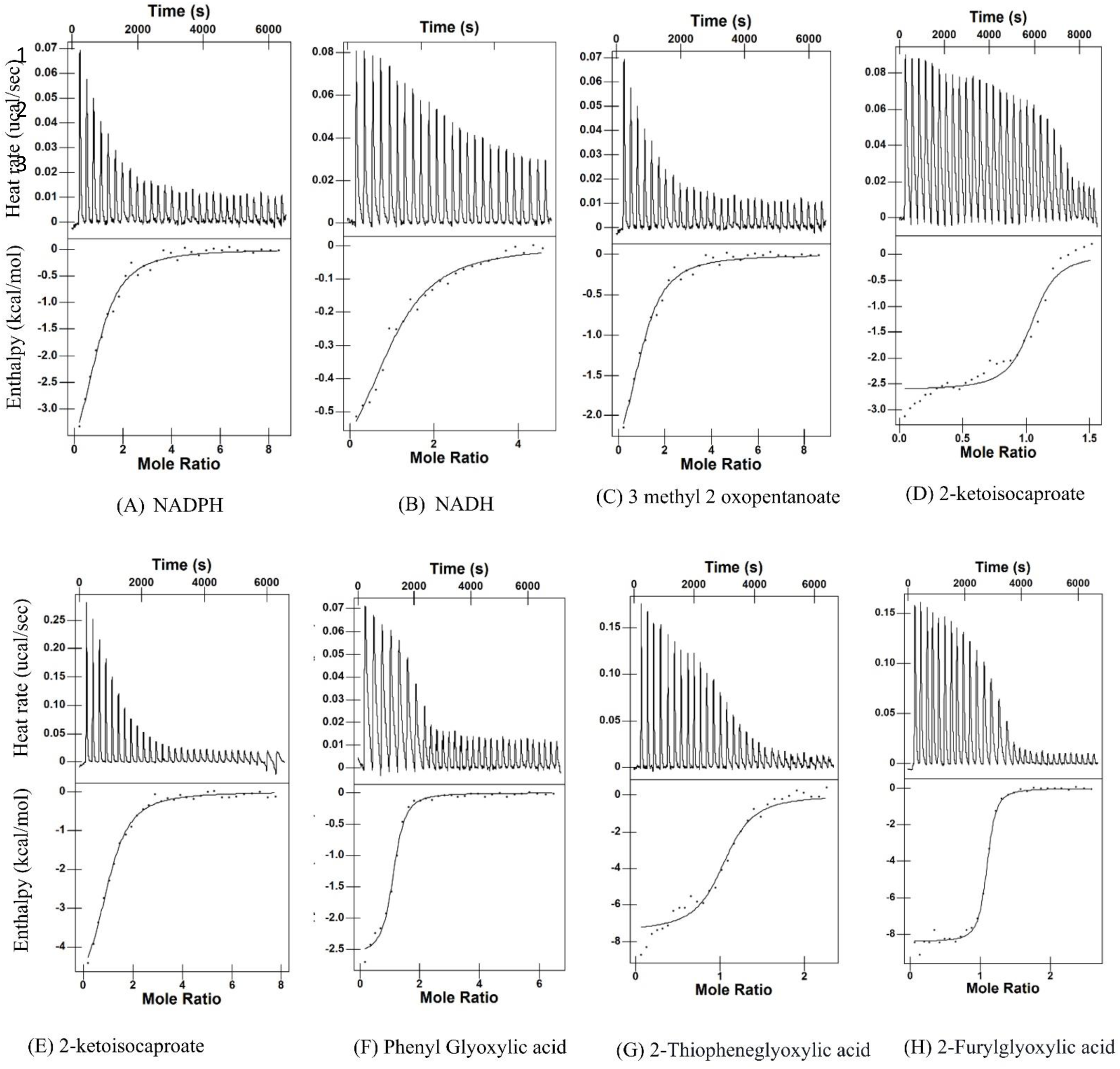
Representative 1TC isotherms for the binding of PaKPR2 to the different ligands (A)NADPH (B) NADH, (C) 3 methyl 2 Oxopentanoate (D) 2-ketoisocaproate, (E) 2-ketoisocaproate, (F) Phenyl Glyoxylic acid, (G) ) 2-Thiopheneglyoxylic acid, (H) 2-Furylglyoxylic acid The lower panels show change in molar heat is expressed as a function of molar ratio of Protein to ligand. In the upper panels, raw ITC data expressed as change in thermal power with respect to time over the period of titration. The solid lines in the lower panels show the fit of data to an independent-site binding model using the integrated NanoAnalyze software as described previously (Wiseman et al., 1989; Mikles et al., 2013).

The residues inside the cleft are mainly active site residues and cofactor binding residues. Cleft closure of the protein enables itself to restrict from solvent exposure. Those conserved residues responsible for cofactor binding have been also highlighted in the MSA results (Supplementary figure 2) with blue colour along with the GXGGX motif highlighted with green colour. From evolutionary perspective the molecular gate present at the surface has never been reported earlier in any of the structure. Finding its role will be interesting as an element of enzyme function evolution. The limitations on the evolution of residues are multifaceted, encompassing a significant influence from solvent exposure. In addition to that, various structural characteristics of the surrounding environment of the residue, such as the depth at which it is buried in relation to the protein’s surface (Chakravarty and Varadarajan, 1999) and the compactness and regularity of the arrangement of neighbouring residues (Hamlryck, 2005) also displays less pronounced effect on molecular evolution of protein function. To investigate further, structure-function relationship of the protein in the context of evolution more aspects needed to be analysed and if required more structural perspectives need to be examined too.

### 2.2 Molecular interaction properties of PaKPR2 with several Ligands suggests that it’s active site may be functional against substrates other than ketopantoate

Characteristic molecular cleft present in all the homologue KPRs converts to the active site of the protein in a ternary complex. However, we observed that this site is partially closed in PaKPR2 and it becomes a two-way pocket. In our next step of analysis, we consider doing interaction analysis with few alpha-ketoacid substrates and cofactors in Isothermal Titration Calorimetry to check whether with altered conformation, the protein is still capable to accept those closely related molecules. For cofactor interaction we decided to go with NADPH, NADH and CoenzymeA because previously these cofactors have been reported to bind with other homologue KPRs in different organisms (Ciulli, A., 2007, Aikawa, Y., 2016). In addition to that we selected our keto acid substrates to be structurally close with ketopantoate, having keto group at C2 carbon (alpha position) and an extended aliphatic or aromatic chain. Substrates tested in ITC analysis are Ketopantoate (KPL), 3-methyl-2-ketopentanoate/ keto-isoleucine (KIL), alpha-keto-isocaproate (KIC), phenylglyoxylate (PGA), 2-oxo-4-phenylbutanoic acid (OPBA), 2-furylglyoxylic acid (FGA), 2-thiopheneglyoxylic acid (TGA), indole Glyoxylate (IGA), indole pyruvate (IPA). (Supplimentray figure 2). All these molecules except FGA, TGA and OPBA can be found in the different biological metabolic systems or pathways.

We considered our ITC interaction to happen only with the protein PaKPR2 in native state and thermodynamic analysis was done for the formation of the binary complexes to understand the receptivity of the individual chemical space present for molecule binding. Thermodynamic parameters obtained in the ITC like K_d_, ΔH and ΔG helps in perceiving these understandings. K_d_ value implicates the strength of binding of molecule in the binary complex which also correlates with the affinity of the molecule. In the process of protein-complex formation, ΔH is reliant on the following: (a) non-covalent interactions between the protein and the ligand, as well as solvation changes at the binding interface; (b) protein conformational changes that result in structural rearrangement in the immediate vicinity of the active site; (c) protonation/deprotonation of ionizable groups of both the protein and the ligand (Luque. I and Freire. E, 2002) and ΔS is reliant on the following: (a) protein-ligand hydrophobic interactions and the release of solvent molecules from buried surfaces; (b) reduction in the conformational freedom of the respective ligand and the protein; (c) decreases in the translational and rotational degrees of freedom of the two parties (Du, X., et al, 2016). The titration trials yielded parameters that demonstrated a significant change in the sign and magnitude of the above-mentioned thermodynamic parameters.

Cofactor CoenzymeA and few substrates like ketopantoate, indole glyoxylate and indole pyruvate showed no thermodynamic changes in the experiment even with high substrate to protein ratios up to 30 times. So, we consider these molecules to be non-reactive against PaKPR2. But NADPH and NADH both cofactors showed binding isotherm with negative enthalpic change of -6.791 kcal/mol and -0.766 kcal/mol respectively. K_d_ value of binding for NADPH is much lower than NADH (7.12uM compared to 98uM) that indicates cofactor specificity of the protein towards NADPH. Interesting finding with ITC is that, despite of being unable to bind its canonical substrate Ketopantoate the protein finds it suitable to bind few other substrates with moderate to High affinity. Keto-isoleucine (KIL), alpha-ketoisocaproate (KIC), phenylglyoxylate (PGA), 2-oxo-4-phenylbutanoic acid (OPBA), 2-Furylglyoxylic acid (FGA), 2-Thiopheneglyoxylic acid (TGA) all these substrates were observed to show exothermic nature of binding isotherm being all their ΔH value as negative with different K_d_ value (Table 3). is presented here with all the thermodynamic parameters for the individual titrations with different ligands. The data implies that structurally closest ketopantoate substrate keto-isoleucine (KIL) is showing lowest affinity. But the substrates with more planer 3-dimensional structure can be related with the higher affinity for the binding site. Substrates with bulky aromatic group in PGA and OPBA also showed good interaction that suggests the available space is not so squeezed to be fully closed. Interestingly, the substrates TGA and FGA that contains 5 carbon ring with sulphur and oxygen atom in the extended region showed best K_d_ value (1.392 uM and 0.392 uM). It implicates that the charged atom (sulphur and oxygen) of the ring is getting stabilized with positive interaction at the binding site. High negative enthalpic value also justify this claim. This means all these substrates can at least have a interaction site within the protein with varying degree of attraction due to limited molecular space. However, only thermodynamic properties from the ITC experiment does not qualify enough to characterise the binding region. Hence, we decided to find structural perspectives of this binding events through crystallography analysis, so that more accurate knowledge can be gained.

**Table 3:** Thermodynamic parameters of cofactor (NADPH/NADH) and alpha keto acid substrates binding to form binary complexes with *P. aeruginosa* KPR2 at 25°C.

| Cell | Syringe | Kd<br>(uM) | ΔH<br>(kcal/mol) | TΔS<br>(kcal/mol) | ΔG<br>(kcal/mol) |
| --- | --- | --- | --- | --- | --- |
| **PaKPR2** | NADPH | 7.68 | -4.702 | -2.276 | -6.978 |
|  | NADH | 98 | -0.766 | -4.516 | -5.283 |
|  | 3methyl 2 ketopentanoate | 11.21 | -2.989 | -3.765 | -6.754 |
|  | 2-ketoisocaproate | 3.048 | -2.609 | -4.663 | -7.273 |
|  | Phenyl glyoxylate | 3.525 | -2.595 | -4.594 | -7.190 |
|  | 2-Thiophenyl Glyoxylate | 1.344 | -7.408 | -0.603 | -8.010 |
|  | 2-Furyl glyoxylate | 0.392 | -5.419 | -3.028 | -8.447 |
|  | 2-oxo-4-phenylbutanoic acid | 9.09 | -5.336 | -1.444 | -6.771 |

### 2.3 Structural insights from ligand bound binary complex of PaKPR2 suggests that the protein’s active site is capable of ligand binding even after being closed partially

ITC interaction analysis gives us substantial proof that the protein is capable of ligand binding with varying affinity. But the site of binding and their binding pattern cannot be elucidated with only interaction analysis. Hence, we tried to solve the binary complex structure of the protein PaKPR2 with 3-mthyl-2-ketopentanoate or keto-isoleucine and NADPH separately. Ketoisoleucine is also a five carbon compound just like ketopantoate. It differs structurally in a way, that the of missing of a methyl group and the terminal hydroxyl group with compared to ketopantoate makes the molecule a bit smaller. So, it is the structurally closest molecule with decent affinity that showed binding in our previous ITC experiments and that’s why we chose to find out its interaction properties with the protein from structural perspectives. Co-crystallization along with crystal soaking with the substrates were employed to get the protein-substrate cocrystal. And after solving the structure (PDB: 8IXH), it was observed that the molecule is present at the one end of the cleft closer to the molecular gate made up of Phe 132 and Tyr 148. It is to be mentioned here that the partial density of Phe 132 was also captured in the structure in closed conformation. That means the residue is dynamic in nature that may affect the entry and exit of molecules inside the pocket beyond the gate. However, molecular space in that region is very small to accommodate all the keto acid substrates. One video is added (Supplimentary Video file 2) to visualize the 3D space of the interaction. The region is turned into small volume pocket due to the presence of the bulky Tryptophan 199 residue that squeezes the size of the void. From the figure 4A, we can understand the molecular interactions between the protein and substrate 3 methyl 2 ketopentanoate/ Keto-isoleucine (KIL). Keto-isoleucine is stabilized by several hydrogen bonds inside the pocket. Mainly the carboxyl group of the molecule forms 3 hydrogen bonds with surrounding residues. Hydroxyl group of this part create hydrogen bond of length 3 Å with Tyr 263 and the carbonyl oxygen found to take part in forming two hydrogen bonds with Asn 200 and Lys 196 of distance 2.75 Å and 2.95 Å respectively. This same Lys 196 also contribute to form one of the two hydrogen bond (length 2.99 Å) with the 2-keto oxygen of the substrate while the second one is with the ILE 133 having a length of approx. 3 Å. Apart from these 4 strong hydrogen bonds, the 3’methyl group of the substrate also makes a stable *π*-alkyl and *π*-sigma bond that is also responsible for the substrate binding. 3D structure of the substrate binding region with the surrounding residues confirms that the space is very limited for the binding of ketopantoate or any larger molecule, like indole glyoxylate and indole pyruvate. But as we noticed that the keto acid group interaction is much conserved, we can assume that smaller substrates may find it easier to go inside and create stable interaction. Whether the substrate interaction finally results in protein’s activity or not depends on the binding of the cofactor also. Hence our next target was to get the complex structure of paKPR2 with NADPH.

**Figure 4:**
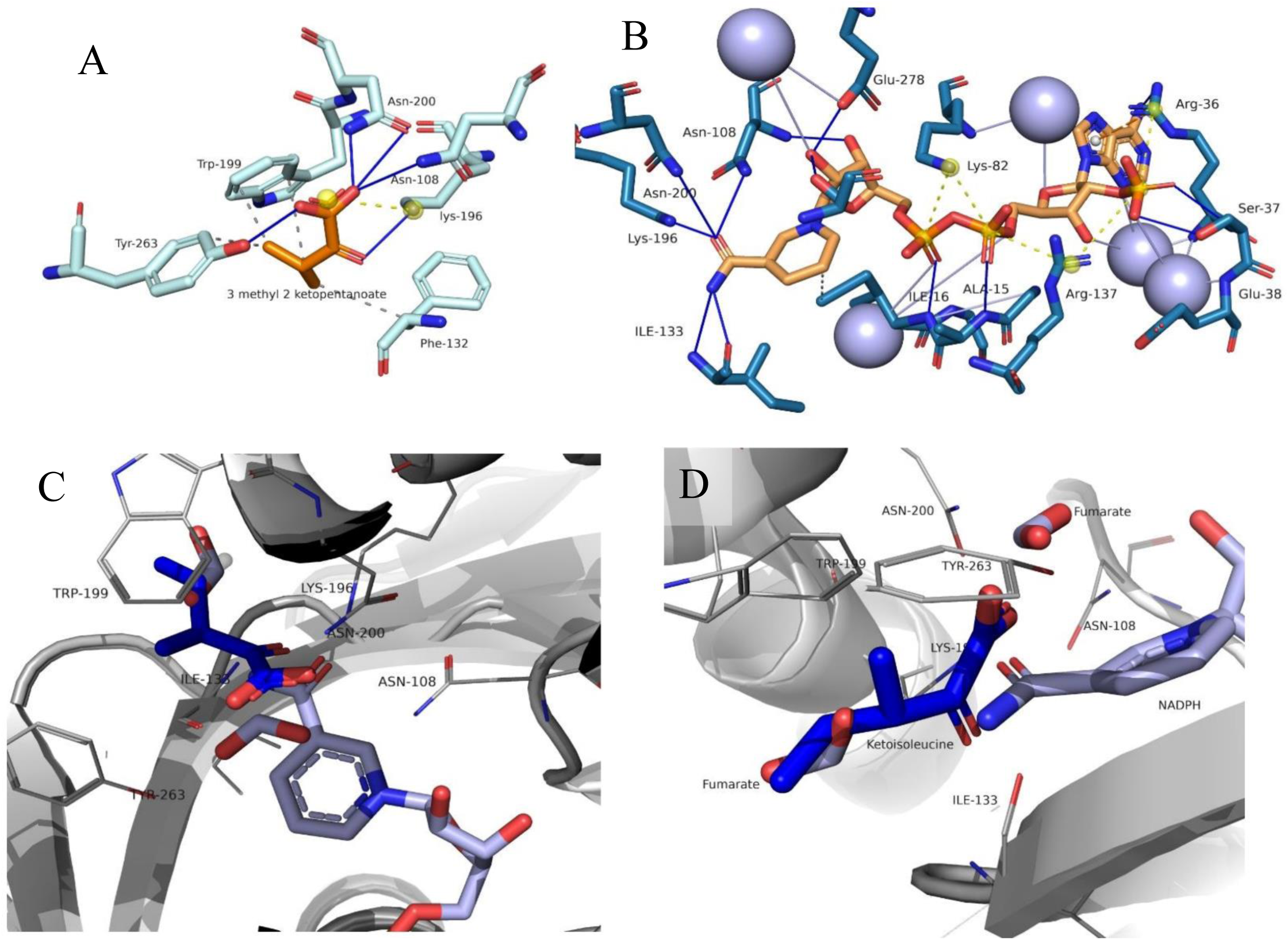
Molecular Interaction profile of Ligands in the Binary complex with PaKPR2. (A) Interaction of Ketoisoleucine / 3methyl2ketopentanoate with the active site residues, (B) Interaction of NADPH with the active site residues.( BLUE lines represent hydrogen bonds and yellow dotted lines represents slat bridges. purple sphere represents water molecules contributing water mediated hydrogen bonds. (C) Relative space sharing by Ketoisoleucine and pyrimidine group of NADPH as analysed by superimposed structures of two binary complexes ( 8IXH and 8IX9).

**Figure 5:**
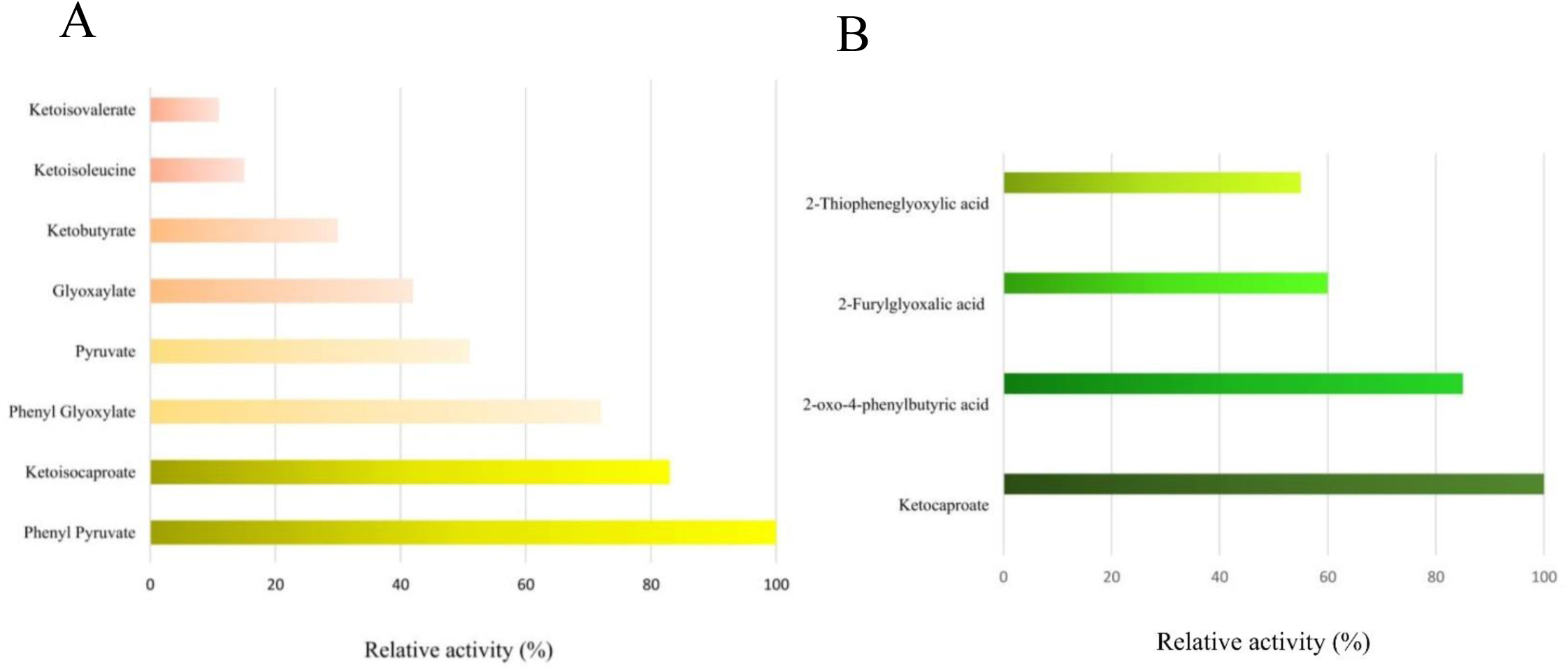
Substrate specificity in terms of relative activity of overexpressed and purified PaKPR2. (A). Specific activity of reductive reaction of different α-Keto-acid substrates (0.5mM) of physiological relevance using NADPH (0.200 mM) as coenzyme. Relative activity was shown as the percentage of the activity of the activity towards Phenyl Pyruvate. (B). Specific activity of reductive reaction of different synthetic α-keto-acid substrates (0.5mM) using NADPH (0.200 mM) as coenzyme. Relative activity was shown as the percentage of the activity of the activity towards keto-caproate.

PaKPR2 and NADPH complex structure was solved with resolution 2.19 and deposited in PDB with ID 8IX9. The protein forms a network of non-covalent interactions – namely hydrophobic interactions, cation-pi interactions, salt bridges and hydrogen bonds to its redox cofactor NADPH. We confirm the cofactor as NADPH (reduced form) from the non-planer electron density of the pyridine ring of the molecule (Supplementary figure 6). Such a reduced form of the cofactor (NADPH) bound binary structure of PanE2/Ketopantoate reductase has not been reported earlier, highlighting a new finding in this study that gives information about the primary stage of the binding which is required for the subsequent activity. Main hydrogen bonded molecular interactions observed in paKPR2-NADPH 3D structure is provided in a video as a supplementary Video file 3. Several dehydrogenases, such as alcohol dehydrogenase, formate dehydrogenase, glutamate dehydrogenase, diaminopimelate dehydrogenase, and the semialdehyde dehydrogenase have all displayed a hinge bending domain closure induced by cofactors (Colonna-cesari, F. et al, 1986: Lamzin, V. S. et al, 1994: Stillman, T. J. et al, 1999; Scapin, G. et al, 1998; Faehnle, C. R. et al, 2006). Even previously reported all KPRs have shown this particular pattern of domain closure upon cofactor binding. But the crystal structure of the KPR2_NADPH_complex compared with apoKPR2 revealed no evidence of a hinge bending induced by the cofactor because the cofactor binding did not result in any significant domain movements in KPR. This was also justified with the comparative volume analysis of the cleft region corresponding to apoKPR and NADPH bound KPR, where no significant change was identified. So, the cleft closure is permanent and not dependent on the presence of the cofactor or any substrate. However, the NADPH binding interaction in the partially closed cleft site is more or less similar to that of other homologues KPR where the complex structure is available. In the crystal asymmetric unit, 3 monomeric unit was present where hydrogen bonds were the main contributor for the interaction in each molecule. Upon further examination, it became evident that specific water-mediated hydrogen bonds and salt bridges are where the true differences lie. The solvent accessible channel that accommodates NADPH found to have 13-16 hydrogen bonds with the cofactor. Figure 4 B shows different moieties of the NADPH forms hydrogen bonds, salt bridges and hydrophobic interactions with the key residues elucidated from the reported structure. Specifically, the adenine group forms 2 water mediated hydrogen bonds and *π*-cation interactions with Arg36, and it fits into a hydrophobic pocket defined by the aliphatic parts of Ile11, Leu34, and Leu92. 2’-phosphate group adjacent to the adenylate moiety interacts with the side chains of Arg36 and Arg137 through salt bridges. The pyrophosphate group is hydrogen-bonded to the backbone amides of Ala15 and Ile16 from the glycine-rich region (GXGXXG). Also, the two extended part of the pyrophosphate found to have carbon hydrogen bond of Gly 80 and Gly 14. Additionally, Lys82 contributes to form salt bridges to this part of the cofactor. The nicotinamide-ribose 2’- and 3’-hydroxyls form hydrogen bonds with Glu278’s carboxylate group, Ser266’s hydroxyl group, and the backbone amide of Asn108. The nicotinamide group also interacts with the side chains of Ile16 through hydrophobic contacts. However, this group exhibits an anti-conformation here, with the terminal group extending outward. It creates three hydrogen bonds with the side chains of Asn 108, Asn200, and Lys196 and two hydrogen bonds with the backbone amide and carbonyl oxygen of Ile 133. This anti-conformation of NADPH at the active site is first time observed among known KPRs. In this anti conformation the Nicotinamide ring of the cofactor is extended towards the substrate binding region and gets stabilized by the bonds created with residues which may also be responsible for the substrate protein interaction. The amino acid W199, along with the nicotinamide ring of NADPH positioned close to the substrate binding site inside the pocket, making it smaller to accommodate ketopantoate or larger molecules, ultimately leading to the enzyme’s inactivity.

So, these two binary complex structure 8IXH and 8IX9 explains that the cofactor and ligand interaction site is exactly at the cleft which is partially closed from one end. However, the molecule binding region is still the same like other homologue KPRs. In addition to the aforementioned findings, the NADPH-bound structure also provides valuable insights into potential substrate bindings. Despite its inactivity against ketopantoate, the protein’s active site may still retain some interaction capability. To gain more knowledge about the active site properties we superimposed these two structures 8IXH and 8IX9 and presented in the Figure 4C and 4D as top view and side view respectively. Here we can see that the pyrimidine ring of NADPH and the substrate keto-isoleucine are positioned in such proximity that they can clash if present together. So in a real scenario, there is competition for this space. However, the void space above the nicotinamide ring can accommodate the planer keto-carboxyl group easily, which is evident from the presence of fumarate molecules in that region seen in the NADPH-bound structure, which were part of the crystallisation buffer. The occurrence of fumarate within the pocket is visible in our complex structure 8IX9. Being the smallest acid unit, the presence of fumarate in the pocket suggests that smaller-volume keto acids could potentially serve as substrates for the protein. This observation opens up possibilities for further investigation into the enzyme’s substrate specificity, which may lead to a better understanding of its catalytic activity.

### 2.4 Activity analysis of PaKPR2 confirms that it is functional against several α-keto-acid substrates in the presence of NADPH cofactor only

Organisms’ capacity to adapt to the dynamic conditions of their habitat is essential to ensuring their survival and reproductive success. As a part of this versatile environmental condition, *P. aeruginosa* also depended on the evolutionary effects on the genes to cope (Winstanley, C. et al,2016). The process of enzyme activity is influenced by the ability to adjust to changes in chemical composition within the surrounding environment, which in turn leads to the modification, transfer, and eventual growth of new functionalities. At the metabolic level, adaptability is linked to the capacity of enzymes to develop advantageous functionalities in response to varying chemical circumstances. Several research investigations have shown important methods via which innovation in enzyme activity may have originated. According to earlier research, D-2-hydroxyisocaproate dehydrogenase of Lactococcus lactis, D-mandelate dehydrogenase of Enterococcus faecium, and possibly other putative KPRs are members of a novel family of D-2-hydroxyacid dehydrogenases that is distinct from the well-known D-2-hydroxyisocaproate dehydrogenase family. Nevertheless, the action of these enzymes is entirely dependent on NADH. *E. coli* ketopantoate reductase was subjected to acting upon several keto acids, including pyruvate, R-ketobutyrate, 2-oxohexanoic acid, R-ketoisocaproic acid, benzoylformic acid, phenylpyruvate, R-ketovalerate, and 3 methyl 2keto valerate (3 methyl 2 ketopentanoate), but only a very low amount of activity was registered for the last two substrates (Zheng, R., & Blanchard, J. S., 2003). So, it is possible that *P. aeruginosa’s* KPR xenologue is specialized to exhibit these activities with the cofactors as an evolutionary intermediate.

Our previous ITC experiments and ligand-bound binary structures satisfied the hypothesis of ligand binding to the active site. In the next step of experiments, we focused on finding the range of the protein’s substrate specificity in terms specific activity. Collectively, we employed 14 α-keto acid substrates, which can be classified into two groups. The first group of substrates are mainly human metabolites like glyoxylic acid pyruvic acid, ketobutyrate, ketovalerate, ketoisocaproic acid, 3 methyl 2keto valerate (3 methyl 2 ketopentanoate/ketoisoleucine), ketopantoate, Phenyl glyoxylate, Phenyl pyruvate, Indole glyoxylate, and indole pyruvate. PaKPR2 failed to show activity against ketopantoate, indole glyoxylate, and indole pyruvate but showed low to moderate efficiencies against others. In addition to this, we also employed the special keto acid substrates that have been known as reactants or precursors for several important chemical reactions, like OPBA, FGA, and TGA. Interestingly, the protein showed much better efficiencies against these synthetic molecules. However, NADPH alone has been found to be the hydride donor for all the reduction reactions, but NADH did not show any activity when used as a co-substrate. Efficacy in terms of specific activity of PaKPR2 against the respective substrates and cofactors at a fixed concentration has been calculated and presented as a bar graph in terms of relative activities. We barred our analysis to find out the steady state parameters like V_m_, K_m_, and K_cat_ to avoid overloading with data that does not comply with the essence of our current study.

PaKPR2’s activity status against the above-mentioned substrates suggests that the protein is capable of acting against several ketoacids; however, that depends on the extended groups of the molecule. Also, the groups having more planer structures increase their chances of getting inside the active site, as evident from the activity against OPBA, FGA, and TGA. Larger molecules may find hindrance in entering the site, resulting in an unsuccessful catalytic conversion that is observed with molecules like ketopantoate, indole glyoxylate, and indole pyruvate. Structural rearrangements at the active site that result in a positive catalytic outcome cannot be judged successfully only from the reaction kinetics or from the structures we have analysed till now. However, a more realistic search for the residue microenvironment at the active site during the enzyme substrate reaction can be achieved by solving the ternary complex structure. This will help us understand if there has been any change in the mechanism of this newly evolved protein and how the changes have affected its efficiency to make it more versatile as a ketoacid reductase.

### 2.5 Ternary complex of PaKPR2-NADP+-KIC elucidate the mechanism of action of the protein and also gives a hint for its moderate efficacy

The preceding sections have thoroughly examined the structural limitations and the residue microenvironment that contribute to the alterations in the functional state of PaKPR2. These findings have also demonstrated its capacity to evolve as a versatile ketoacid reductase. The activity status, as determined by several substrates, indicates that it is not exclusively confined to the substrates listed here. The protein PaKPR2 has the ability to convert additional substrates, and the underlying reasons for this have been clarified by a thorough analysis of its structural features. We expanded our study to analyse the extent of limitations that the protein is encountering from a mechanistic standpoint due to the changes caused by partially closed ceft. The ternary complex of Ketoisoacproate-NADP-PaKPR2 structure will surely add in more information about the real time perspectives of activity and intricacies of the enzyme function. Considering this we set up co-crystallization trials with NADPH and the substrate Keto-isocaproate (KIC). In our initial crystal structures, KIC electron density was not good enough due to lose binding hence we soaked our crystals in 5mM KIC containing cryo buffer for 15 mins before data collection. Finally, after repetitive data collection, in the solved ternary complex structure that was deposited in PDB with ID 8IXM. a prominent density of the substrate and Cofactor was visible in the electron density map. 2Fo-Fc omit map of the densities has been provided in the Supplementary figure 6.

The first striking observation of this structure (8IXM) reveals a shift in the orientation of the nicotinamide ring of NADPH, in comparison to the NADPH bound binary structure (8IX9). The planarity of the group indicates that it is in the oxidized state as NADP+. We have mentioned earlier that the extended conformation that we observed in NADPH bound structure is not reported earlier in any of the previously reported structure. However, the orientation of the cofactor after the rotation of the pyrimidine ring by 90° (Figure 6A)., observed in the ternary complex is more similar to the conformation of other cofactors found in various KPR-cofactor binary complexes reported in the Protein Data Bank (PDB :5HWS, 4H3M etc.). Regarding the interaction of the NADP+ within the active site pocket, there have been few differences found, specifically in two opposite ends of the molecule. First, the rotation of the nicotinamide ring results in the formation of hydrogen bonds between Tyr 263 and Val 135 with the group. In the opposite end, it is seen that the adenylate group forms an additional hydrogen bond with either Glu 88 or Gly 85. Actually, in two molecules of asymmetric unit in the structure recorded the difference in movement of the adenylate group (Figure6B). Therefore, in one chain, the adenosine moiety forms a hydrogen bond with the side chain of Glutamate 88, whereas in another chain, the bond is formed between the moiety and the amide group of the glycine 85 residue. This extra interaction implies that during catalysis NADPH needs more stability.

**Figure 6:**
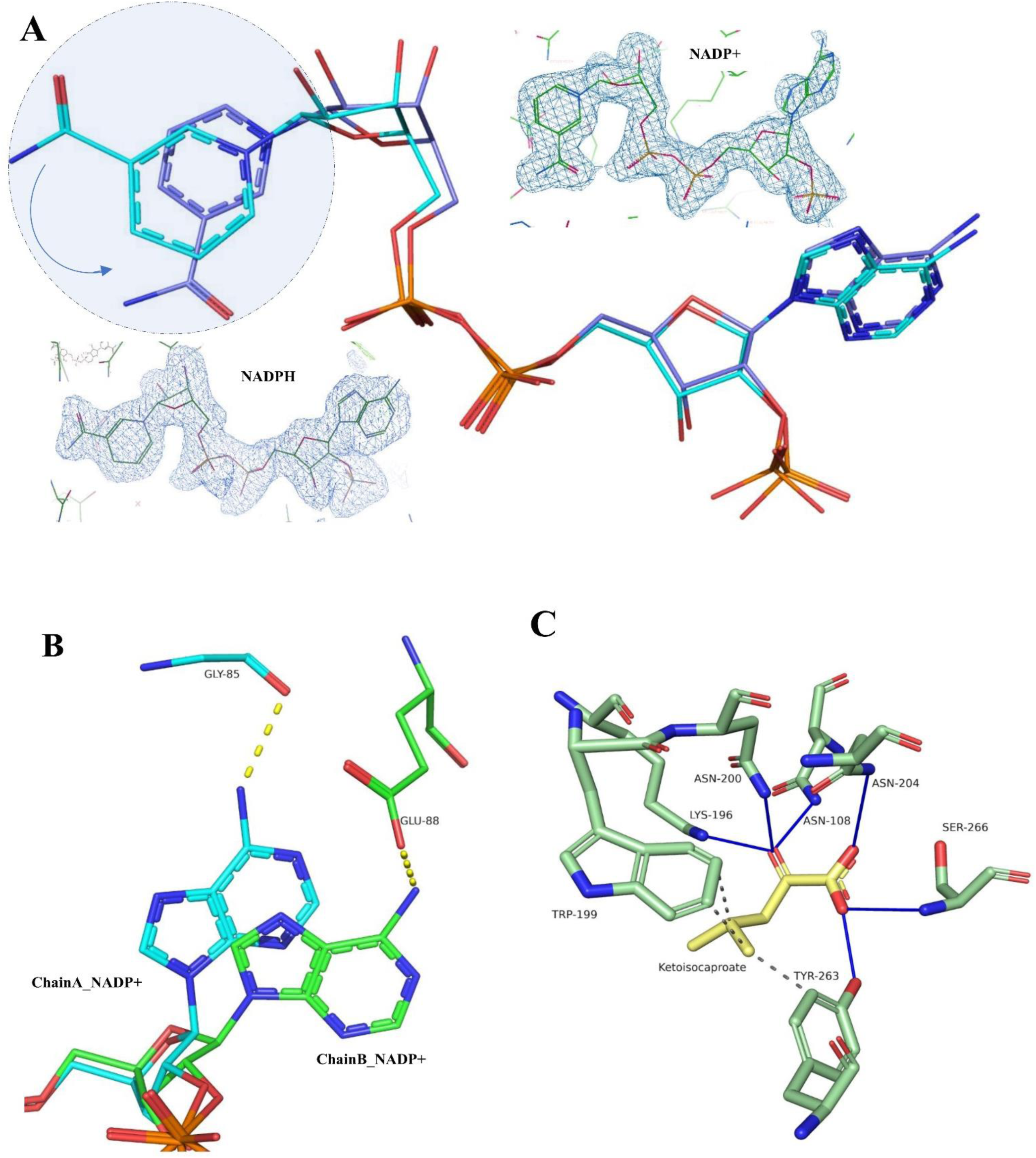
Ligand interactions in Ternary complex of PaKPR2-NADP+-KIC. (A) Relative orientation change of Nicotinamide ring of NADPH in ternary complex (purple) as compared to that in binary complex (cyan colour)-Highlighted region shows the rotational movement, directed with an arrow, respective 2FoFc maps of the cofactors in two different structures are in inset pictures. (B) Extra hydrogen bond observed between adenylate group of the cofactor in ternary complex with two different residues in two different chains of the asymmetric unit. (C) Interaction of ketoisocaproate with the active site residues in ternary complex.

Substrate KIC and Protein PaKPR2 interactions observed in the ternary complex is really fascinating and also justify our initial hypothesis regarding fumarate binding in the NADPH-KPR binary complex. First, the placement of the molecule is much deep inside the pocket than the substrate in binary complex (Ketoisoleucine in 8XIM). So, there is a change in the interactions that stabilized KIC inside the pocket. Asn 108, ASn 204, Lys 196, Tyr 263, Ser 266 residues forms strong hydrogen bonds with the molecule. Due to these interactions the functional group of KIC is placed just above the nicotinamide ring at a distance less than 4 Å. This distance is very much important for hydride (H+) transfer during enzyme function. All productive electron transfer reactions in biology fall within a distance range of 4 to 14Å. (Moser, C. C. et al, 2010). Most importantly, the existence of a significant hydrogen bond involving the lysine 196 residue, which is essential for substrate reduction, is the primary indication that our structure is a productive complex that can be seen in our ternary complex structure. This contact is established as a conserved interaction in all other ternary complexes that have been previously reported.

If we consider previous reaction scheme described for *E.coli* KPR (Ciulli, A. et al, 2007) general mechanism of substrate reduction step includes the transferring of C4 *pro-S* hydride of NADPH to the *si* face of keto-acid group, with protonation of the developing alkoxide by the hydrogen bonded Lys 196 residue. We assume, the orientation of the cofactor’s ribose-nicotinamide moiety and the placement of the substrate nearest to the moiety is necessary for efficient hydride transfer and fruitful substrate binding. The carboxylate group of the substrate forms hydrogen bonds with the conserved Ser266, locking it in place at each stage of the reaction. The side chains of four residues—Asn108, Asn200, Tyr 263, and Trp 199—provide additional binding interactions. The loss of hydrogen bonds from Lys 196 and Asn 200 facilitates the release of substrate from the ternary complex. The catalytic cycle is terminated by the rate-limiting release of NADP+. The alkoxide formation of Lys 196 during catalysis initiation and re-protonation of the residue during product release depends on the solvent molecule, that is also observed in the structure as a presence of a water molecule in the vicinity of the residue Lys 196.

Ther ternary complex guided us to describe the scheme of mechanism for PaKPR2 for the NADPH dependent reduction activity. And the residues involved in the reaction found to be conserved with respect to Ketopantoate reductase of homologue origins. However, the main difference here is during the function no hinge movement is present. All the reaction is occurring inside the two-sided pocket. It is noteworthy to mention here that, in different structures Tyr 263 and valine 135 molecule was involved in forming hydrogen bonds either with each other for the cleft closure or with water molecules. Here in ternary complex also these two residues have been found to make hydrogen bonds with the substrate and cofactor. Hence it can’t be an overstatement that these residues present in two opposite domains are actively contribute in maintain different conformations of the protein. Specifically, Tyr 263 present in a loop regain is very dynamic during molecular interaction and subsequent activity. These observations also suggests that a very small re-orientation of the active stie takes place before or during the catalytic events.

## Discussions

In a broad perspective the residues present and controlling the functional adaptability of a protein are of real significance on evolutionary perspectives. Protein evolution study through sequence and genomic content analysis is widely accepted and those attempts brought an abundance of valued information related to the field. But visualizing those well accepted hypothesis is always been a barrier for the evolutionary biologist. The study here bridges the gap between the structure and evolution study by keeping functional attributes as a background protagonist.

On this notion our investigation focuses on the second copy of Ketopantoate Reductase (KPR) in *P. aeruginosa*, namely PaKPR2. Its intriguing inactivity against the natural substrate Ketopantoate despite of maintaining active site reside conservation prompted us to find out its structural and function status. Through several crystal structures and biochemical analysis, we explore the cause behind its evolutionary changes that results in its current functional traits. By exploring the intricate relationships between structure and evolution, our goal is to uncover the functional roles, metabolic specializations, and evolutionary constraints associated with the widespread multiplication of this enzyme across diverse organisms.

Initial finding from the apoKPR2 structures reveal a that the sequence evolution transformed the active site cleft in to a two-sided pocket. The cleft that present in between two globular domain of the protein is actually the site for ligand binding and catalytic functions. This cleft, considered as a characteristic element of this protein, is essential for hinge movement during catalysis in all the homologues. We reported the changes in the residue interaction and motif movements that caused the partial closure of the cleft in PaKPR2. It seems the volume sequestration converted the large open cleft into a pocket with lesser space that primarily caused its inactivity towards its substrate Ketopantoate. Notably, there’s a subtle difference in the two native structures reported here. In one, the pocket is closed at one end, while in the other, it has an opening, forming a two-way pocket. This pocket is a probable substrate binding site too. We also reported the dynamic movement of residue F132, interacting with Y148, creates a molecular gate at this end. The open conformation and the available space at the site imply that substrates with smaller size may go inside and interact with the conserved residues while the lesser space was also a concern for cofactor binding.

Despite cleft closure, molecular interaction properties and activity analyses suggest that PaKPR2 exhibits versatility as a ketoacid reductase. ITC and activity assay with 14 different Keto-acid substrates showed here confirms the above-mentioned claims. However, the protein confers variable affinity towards those substrates indicated by the thermodynamic as well as the kinetic parameters. The limited space for binding and the molecular regulation at the entry exit site (presence of molecular gate) may cause the protein to be a hysteric in nature of function. It is to important to mention here that the protein showed significant rate of function against few synthetic keto-acid molecules like FGA and TGA that suggests its capacity as the biocatalyst for the reductive conversion of those molecules. Newly evolved protein can serve as a more potent biocatalyst as well as studying their evolutionary path can guide us to elucidate the unknown features of the natural episodes of alteration and modifications.

In addition to that, the claim that the protein is exerting different constraints through residue microenvironment at the binding sites, it was necessary to justify by visualizing the structures with ligand and cofactors separately. Hence, the structures of binary complexes with one of the substrates (Ketoisoleucine) and NADPH cofactor were solved that represents the conserved and restricted ligand-binding interactions at the active site. It also clarifies that the space available at the active site is the actual cause of proteins inability to accept all the keto acids molecule as substrate and the activity also depends on the molecules’ appropriate shape and size for accommodation.

Finally, PaKPR2’s functional evolution as a versatile keto acid reductase was thoroughly investigated by analysing a ternary complex structure, PaKPR2-NADP+-KIC. How the protein having a limited space of ligand binding is still maintaining its functional status has been visualized in this structure. Molecular arrangements during catalytic events were noted, and based on that, a scheme for controlled mechanisms was interpreted. This study explains the roles of conserved residues and other auxiliary residues in maintaining the protein’s functional status.

The results presented in our manuscript provide detailed visual experiences of various structural perspectives in the molecular evolution of PaKPR2. We believe that the empirical nature of our study contributes substantively to the understanding of protein evolution in real-world systems. Over billions of years since the Last Universal Common Ancestor (LUCA), duplicated genes have led to diverse enzyme superfamilies. Initially, generalist enzymes evolved into specialists, and later, promiscuous activities from specialized enzymes diversified their functions. Elevating promiscuous activities to physiological relevance is complex, influenced by environmental conditions, available activities, metabolic network topology, and the enzyme’s original and novel function. Hence, the present status of the PaKPR2 as a nonfunctional enzyme against ketopantoate doesn’t make it totally dead. However, nature tried to restrict its function through several structural constraints and helped in sustaining its place in the genome as an evolutionary intermediate waiting for its turn to become more potent with fixed and specific role in the organism.

## Materials and methods

### Bioinformatic analysis and Multiple sequence alignment

*Pseudomonas aeruginosa* ketopantoate reductase gene sequence was retained from the KEGG database (Kanehisa.M, 2000). Structures of KPR homologues and their respective sequences were retrieved from the Protein Data Bank (PDB). Sequence similarity was analysed by pBLAST server (Altschul, S.F., 1990). Multiple sequence alignment was done with the sequences in MEGA11 software package. (MEGA11: Molecular Evolutionary Genetics Analysis version 11, Tamura, Stecher, and Kumar 2021.)

### Cloning, expression, and purification of ketopantoate reductase

The second copy of panE gene in P. aeruginosa UCBA PA-14 strain with NCBI-Protein ID: <u>ABJ10938</u> , was inserted into the Nde1/Sal1 restriction enzyme site of the pET28a (+) expression plasmid vector and introduced into E. coli BL21 DE3. Primary cultures were grown overnight in LB medium for inoculation in small volume (10ml), which was added in the secondary cultures of 600ml. It was left under shaking at 37°C, until reaching an optical density (OD) of 0.6 at 600nm. Subsequently, the recombinant protein was expressed by 1mM IPTG induction for 4 hours at 37°C. Post-induction, cells were harvested and the pellets were re-suspended in lysis buffer containing 20 mM Tris pH 8.0, 150 mM NaCl, 10% glycerol, 1 mM PMSF and 5mM Imidazole. After sonication and centrifugation at 20000g for 40 minutes, the cell lysate was transferred to a pre-packed Ni-NTA agarose beads column which was pre-equilibrated with equilibration buffer. Beads were then washed with a wash buffer, and the protein was eluted using an elution buffer. The equilibration, wash, and elution buffers shared a composition of 20 mM Tris pH 8.0, 150 mM NaCl, and 10% glycerol. Additionally, the equilibration, wash, and elution buffers contained 20, 40, and 250mM imidazole, respectively. The eluted protein underwent column chromatography using a Superdex HiLoad 16-600 gel filtration column. The protein was concentrated using a Millipore Amicon Ultra centrifugal column with a 30kDa cut-off. The filtered and concentrated samples were then employed for crystallization.

### Crystallization

The concentrated and filtered samples were utilised for the process of crystallisation. The Rigaku wizard classic screen was used to primarily screen crystallisation conditions. TTP Labtech MOSQUITO robotics unit was used for setting up the sitting drop MRC SD2 plates for primary screening of the crystallization conditions. The growth of crystals under favourable conditions was seen in the screening plate, and subsequently, these conditions were optimized utilising the TTP Labtech Dragonfly robot unit. Crystals of native Protein grew after 24 hours in the condition E3 of wizard classic 2 that contains 20%PEG 8000, 100mM Tris-HCl pH 8.5, 200mM MgCl_2_. Binary complex of paKPR2-Ketoisoleucine was optimized with the condition 11% PEG 6000, 100mM HEPES pH 7.5, 200mM MgCl_2_,substrtes were added during co crystallization and before concentrating the Gel filtration elute. PaKPR2-NADPH complex structure was attained by co crystallization of the substrate and protein in the screen condition A4 0f Wizard classic 3. There was no need for optimization for this crystal as the crystal in the screening plate was well enough to get the final data. Finally, for the ternary complex preparation diluted protein PaKPR2 of concentration 1mg/ml, 5mM substrate Ketoisocaproate and 1mM cofactor NADPH were mixed together and left for overnight in 4° C. Crystals were obtained in the screen condition of 16% PEG 8000, 100mM CHES 9.5. Rigorous optimization trials were needed for the final data collection as we were getting partial substrate density. For successful crystal we added 1 to 2mM of substrates in a series of increasing concentration as a part of the cryo solution.

### Data collection, structure solving and refinement

All the diffraction data except the binary complex with NADPH were acquired at a temperature of 100 K using the in-house Bruker D8Ventura machine equipped with a Photon III plus detector.PaKPR2-NADPH crystal were diffracted in INDUS-2 Beamline PX-BL21 at RRCAT, Indore (Department of Atomic Energy, Govt. of India) and the data was collected on MAR detectors. Irrespective of diffraction source, prior to mounting the crystals under a nitrogen air stream, each crystallization solution, enriched with 10-15% glycerol, served as a cryoprotectant. For successful complex crystal data collection substrates were also added during cryo-soaking as and when needed. The obtained data underwent indexing, integration, and scaling using PROTEUM3 software that comes with the machine. Two PaKPR2 native data were solved in the P 1 21 1 space group, with 2 monomers in the asymmetric unit. They were deposited in the PDB with ID 8IWG (resolution 2.15Å) and 8IWQ (resolution 2.19Å). In addition to that the binary structures were solved in the C 1 21 space group, with 3 monomer containg asymmetric unit. They were deposited with PDB ID 8IX9 (resolution 2.20 Å) and 8IXH (resolution 1.96Å) having NADPH and Ketoisoleucice as ligand respectively. The ternary complex structure belongs to the P 1 2 1 space group, featuring a 2 monomer chains in the unit cell. The resolution was 1.96 Å and it was deposited in PDB with ID 8IXM. The overall quality of the X-ray data was assessed using PHENIX.xtriage (Adams, P.D. et al, 2010). Molecular replacement was done with the PHENIX.phaser (Mccoy, A.J. et al, 2007) module was used to solve the apoKPR structures, employing single chain of the the native panE2 structure with PDB code 1KS9 as a template model. Subsequently, all structures were solved using our native structure 8IWG_chainA as a model. The initial model for each structure was manually built in COOT (Emsley, P. et al, 2004) and refined iteratively using PHENIX.refine (Adams, P.D. et al, 2010). Ligand geometric restraints were generated using eLBOW (Moriarty, N.W. et al, 2009). The resultant models underwent verification for geometric and stereochemical quality using MolProbity software (Chen,V.B. et al, 2010) All structures were deposited in the PDB, and the respective validation reports were obtained. Visualization of structural properties and the generation of molecular images were conducted using Chimera (Pettersen. E.F., 2004), PyMol, or Discovery Studio Visualizer.

### Computational volume analysis of solvent accessible channels and pockets

In order to measure the solvent accessible volumes and corresponding surface area of cleft regions in our deposited structures as well as previously reported KPR sturctures ,CASTp web server (Dundas, J, 2006) were utilised. The service provided surface representations of void areas with measurable volumes. In Chimaera visualisation software (Pettersen. E.F., 2004), the volumes of the ligand binding pocket and cleft were visualized for analysis.

### Isothermal Titration Calorimetry Analysis

Using a TA Nano-ITC device, isothermal titration calorimetry (ITC) tests were carried out. Briefly before experiments, wildtype PaKPR2 and the ligands (Supplementary figure 3) were dialyzed individually in the ITC assay buffer containing 20 mM HEPES ph 7.5, 50mM NaCl and 2% Glycerol. All experiments were initiated by injecting 30 × 3-μl aliquots of 200–800 μM of Substrates and cofactors from the syringe into the calorimetric cell containing approx. 200 ul of 30-50 μM of the PaKPR2 protein at 25 °C. Integrated NanoAnalyze software that comes with the instrument automatically recorded the change in thermal power as a function of each injection. After further integration of the raw data, binding isotherms of heat release per injection as a function of the substrates to PaKPR2 molar ratio were obtained. Titrations of the ligand to the buffer were carried out in order to accommodate baseline adjustments. After that, the adjusted heat change was fitted, using the nonlinear least squares regression analysis present in integrated NanoAnalyze programme. The binding isotherms were iteratively fitted to a built-in Independent binding model as previously mentioned (Wiseman et al., 1989; Mikles et al., 2013). , yielding the stoichiometry, **n**, the enthalpy of binding, **ΔH** , and the dissociation constants, **K_d_**. Curve-fitting of data acquired for Substrates with low c values was fixed at a constant stoichiometry of 1, which was established from titrations with NADP(H) under high c values circumstances. The equation ΔG = RT lnKd (1), where R is the universal molar gas constant (1.99 cal/mol/K) and T is the absolute temperature (298 K), was used to compute the free energy change (ΔG) upon binding. With ΔH and ΔG, the entropic contribution (TΔS) to the free energy of binding was computed using the following formula: TΔS = ΔH - ΔG (2). All calculations were automatically provided by the NanoAnalyze software with successful graph fitting.

### Enzyme kinetics

The standard assay mixture contained 25 mM sodium phosphate buffer (pH 7.0), 0.5 mM of different α-keto-acid substrates, 0.200 mM NADPH, and sufficient amount of enzyme in a final volume of 3.0 mL. The reaction was initiated by the addition of substrate, and the decrease in absorbance at 340 nm (A340) was measured at 35°C, which was performed using an UNICO 2802 UV/VIS spectrophotometer. A molar extinction coefficient of 6.22 cm2/pmol NADPH was used for the calculation of enzyme activity. One unit was defined as the amount of enzyme caused the oxidation of 1 μmol of NADPH per min. Maximum rate of reaction was noted for each substrate and the relative activity was measured according to that. Substrates were classified in 2 categories, one having physiological significance and another one with synthetic relevance. For the first category Phenyl pyruvate was showing maximum activity while for the other category it was ketocaproate.

## Supporting information

Suppliment figue and table

## Data availability

All data are contained in the article or available on request by contacting the corresponding author

## Supporting information

This article contains supporting information.

## Acknowledgements

This work was partly supported by grants from CSIR-IICB (P-07). GBC acknowledges Department of Biotechnology (DBT) for fellowships. We thank Dr. Ravindra Makde and his team for providing an opportunity to collect X-ray diffraction data at PX-BL21, INDUS-2, RRCAT, Department of Atomic Energy, and Government of India. GBC also thank Dr. Basavraj Khanppnavar and Dr. Abhisek Mondol for giving guidance and training as lab seniors. Also, very thankful to Dr, Rajeev Kumar and Dr. Chittran Roy for their valuable suggestions during the work.

## Author contributions

SD and GBC conceived the idea and designed experiments. GBC carried out all molecular biology and biochemical analysis; performed protein crystallization, structure solving GBC And SD did Structural analysis; SD supervised in all the steps. GBC wrote the manuscript; SD reviewed all the data and the manuscript. All the authors gave editorial input.

## Abbreviations

The abbreviations used are

KPR: Ketopantoate reductase
NADPH: Nicotinamide Adenine di Nucleotide phosphate hydrogen
KIC: Ketoisocaproate
KCA: Ketocaproate
KIL: Ketoisoleucine
HEPES: 4-(2-hydroxyethyl)-1-piperazineethanesulfonic acid
CHES: N-Cyclohexyl-2-aminoethanesulfonic acid

