## Supplementary material for "Implication of structural constrains facilitating the functional evolution of *Pseudomonas aeruginosa* KPR2 into a versatile α-keto acid reductase": Suppliment figue and table

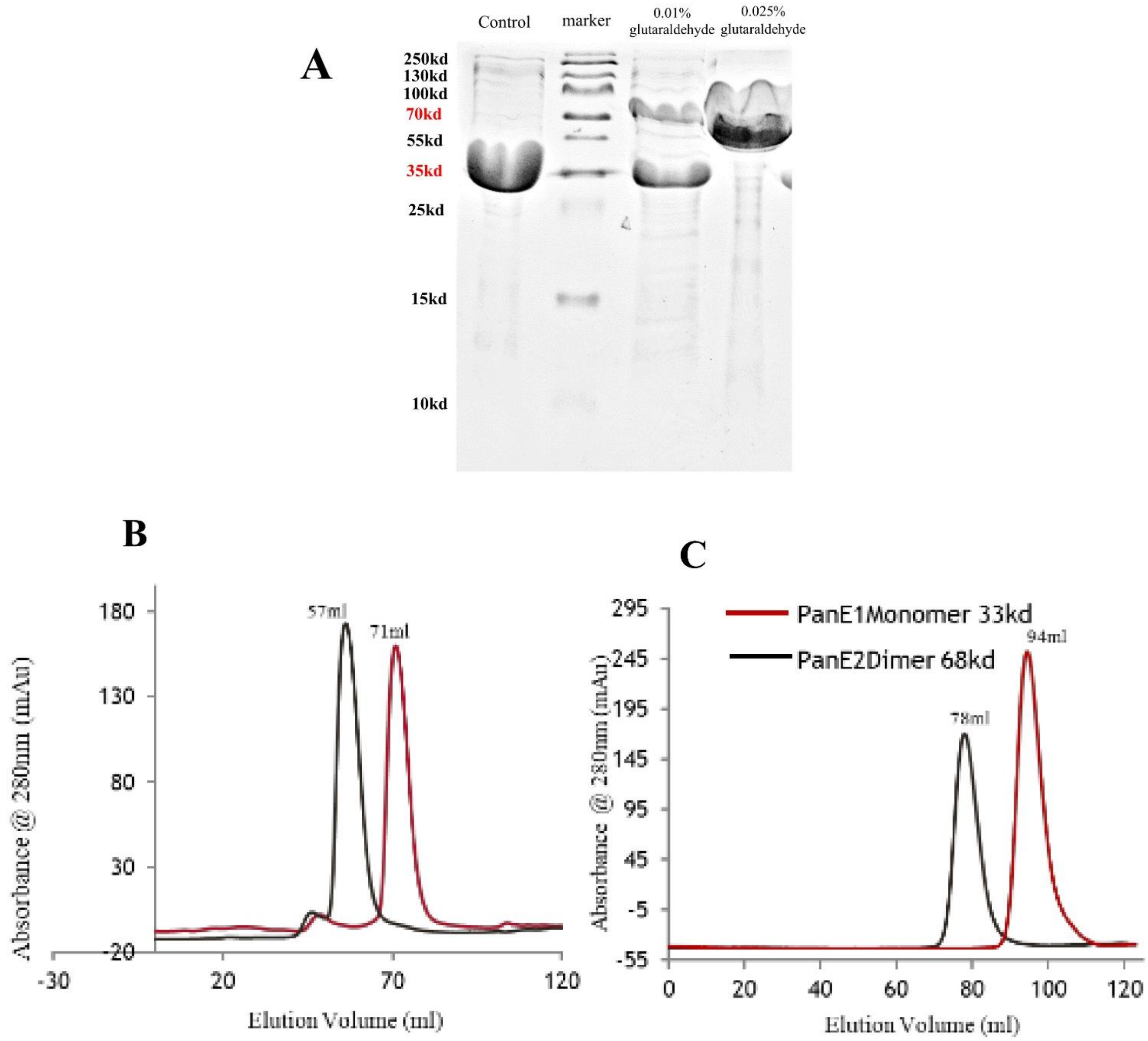


**Supplementary Figure 1: Analysis of oligomeric state of PaKPR2** (A) Gluteraldehyde Crosslinking of PaKPR2 (B,C) Gel filtration profile of PaKPR1 and PaKPR2 with two different Gel filtration column having different resolution range.

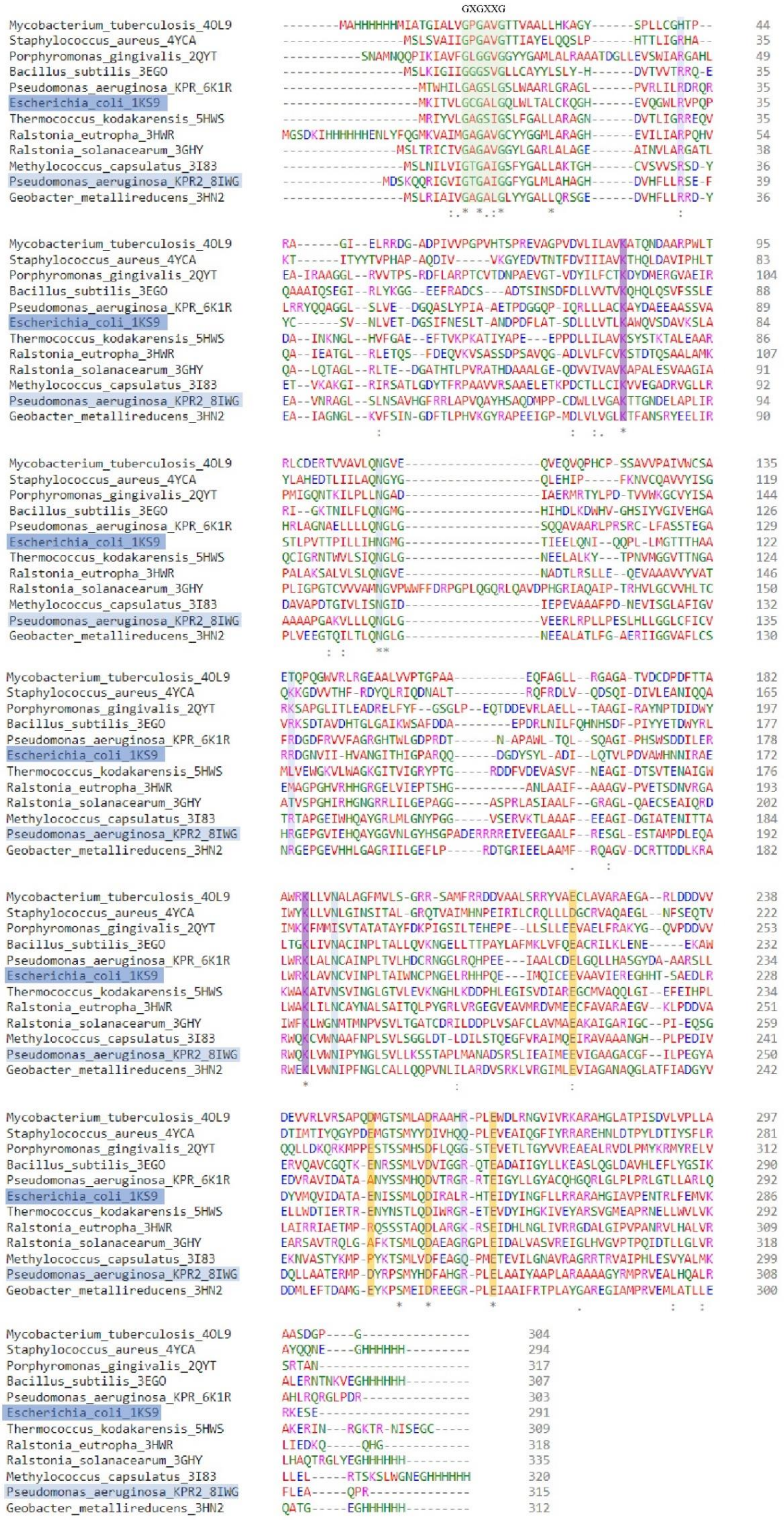


**Supplementary Figure 2: Multiple sequence alignment of KPR homologs those have structures in PDB**

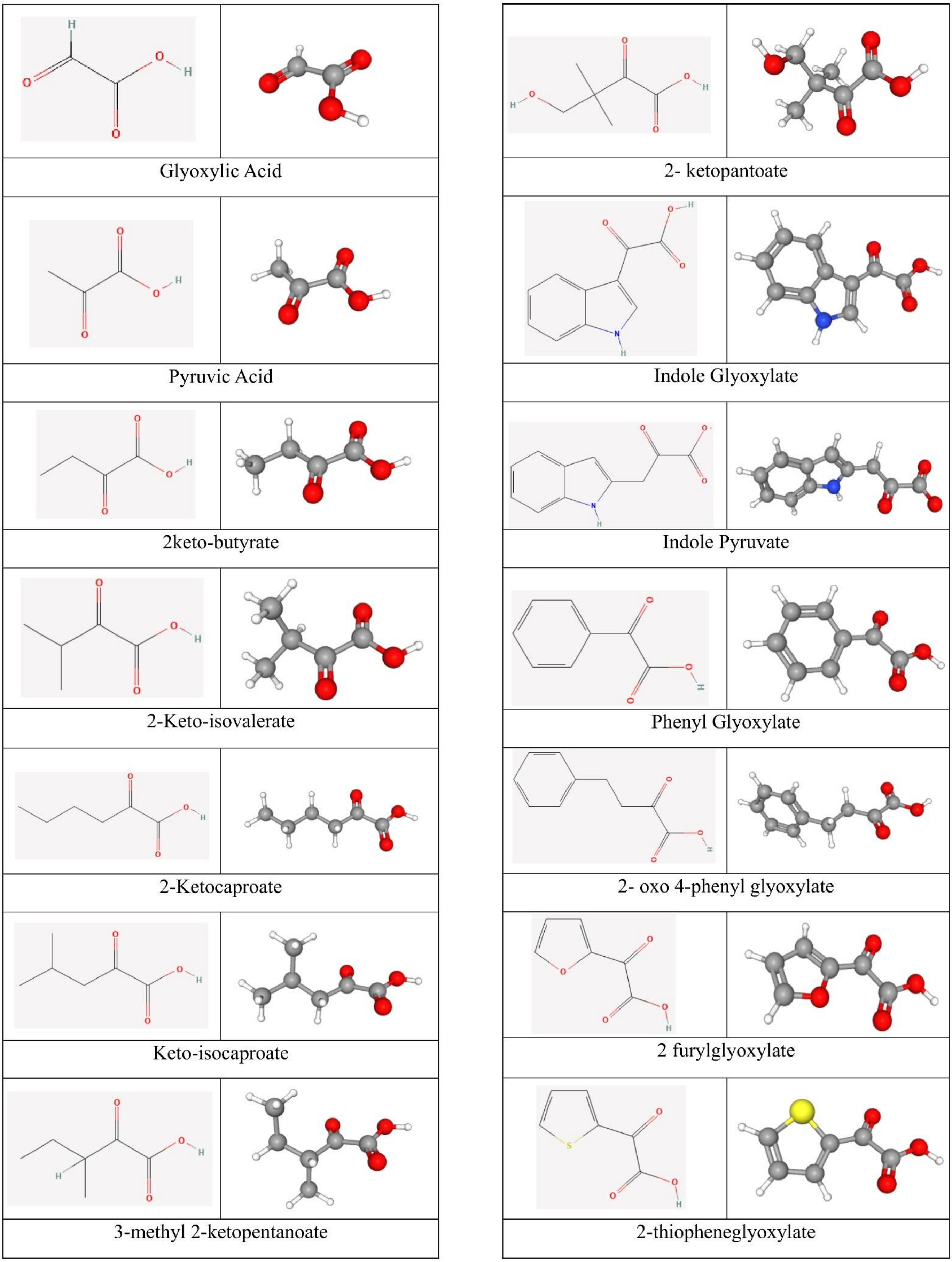


**Supplementary Figure 3: Various α-keto acid structures in 2D and 3D form.**

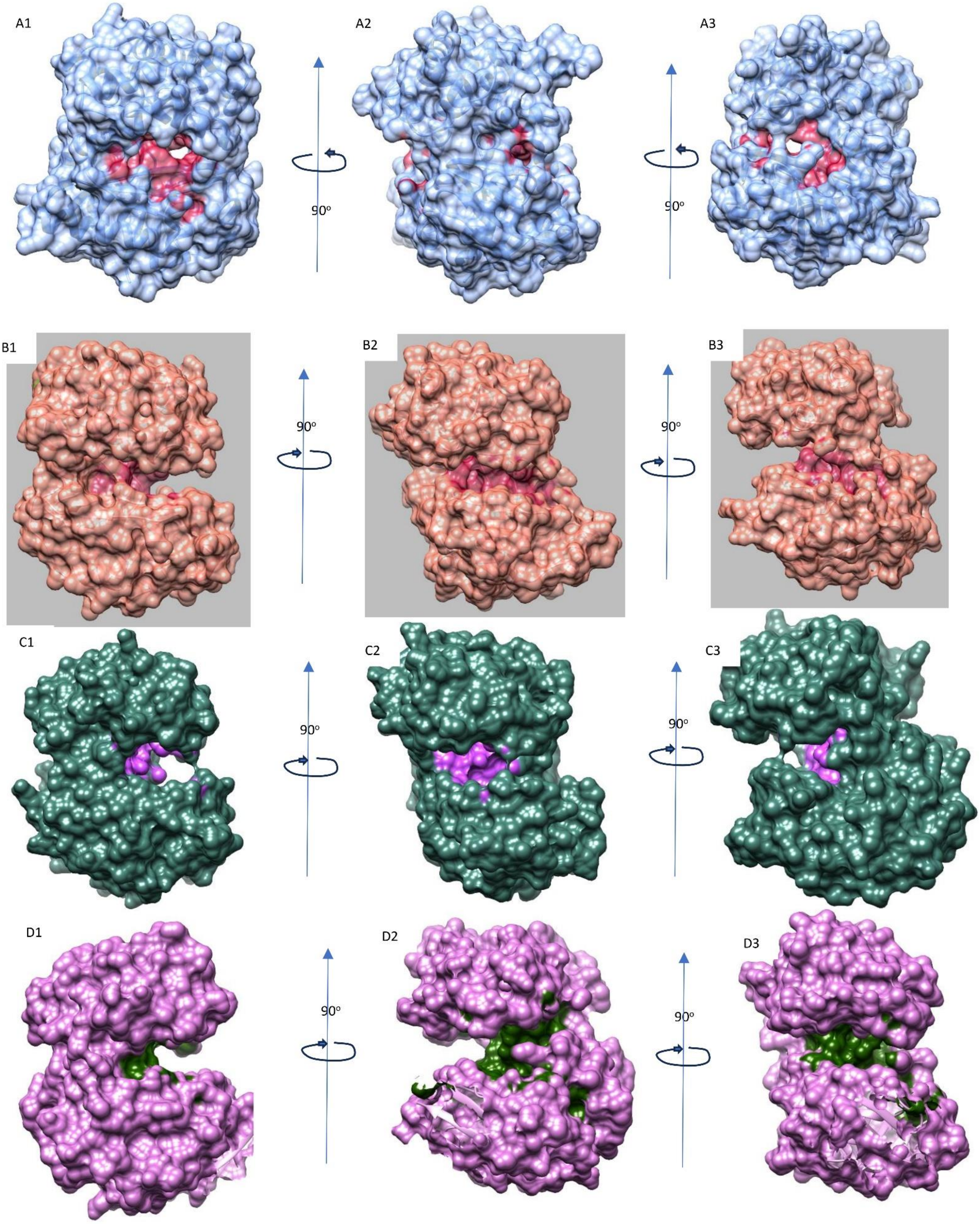


**Supplementary Figure 4 – Surface representation of three 90º apart views of the cleft** (A) partially closed Cleft turned Pocket of *Pseudomonas aeruginosa* KPR2 (8IWG) compared to open cleft of apo KPR1 of (B) *Pseudomonas aeruginosa* KPR1 (5ZIX) (C) , *Escherichia Coli KPR* (1KS9) and (D) *Bacillus subtilis* KPR (3EGO)

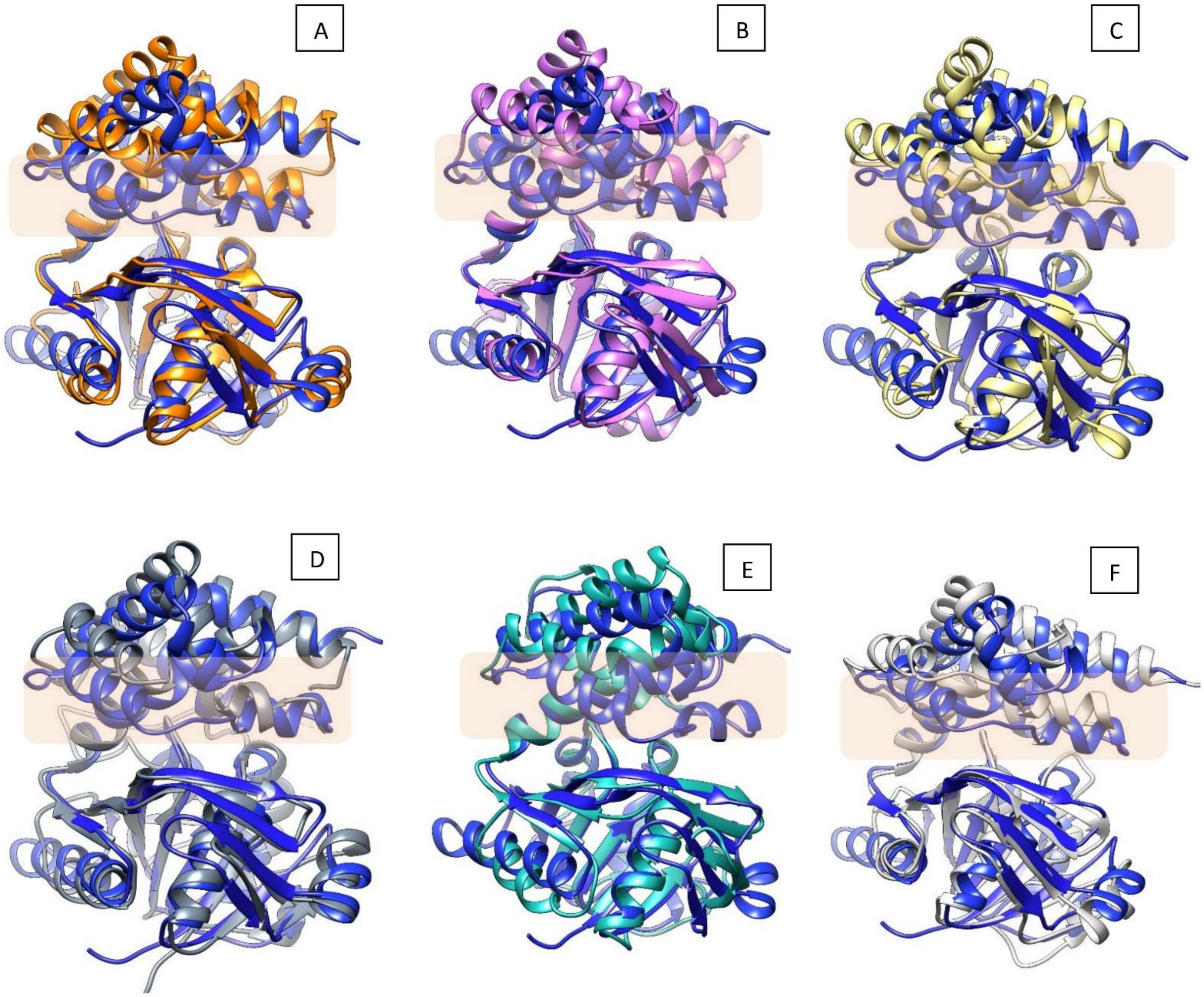


**Supplementary Figure 5 – Superimposition of apo PaKPR2 with other apo KPR structures available in RCSB PDB, Highlighted region of blue structures (paKPR2) showed the HELIX-TURN HELX motif closer to the opposite domain that made the cleft volume smaller.**

(A) *Pseudomonas aeruginosa* KPR2 (8IWG) – deep blue, *Pseudomonas aeruginosa* KPR1 (5ZIX) –orange
(B) *Pseudomonas aeruginosa* KPR2 (8IWG) – deep blue, *Escherichia Coli KPR* (1KS9)- light pink
(C) *Pseudomonas aeruginosa* KPR2 (8IWG) – deep blue, *Bacillus subtilis* KPR (3EGO)-light yellow
(D) *Pseudomonas aeruginosa* KPR2 (8IWG) – deep blue, *Ralstonia solanacaerum* KPR (3GHY)- grey
(E) *Pseudomonas aeruginosa* KPR2 (8IWG) – deep blue *Staphylococcus aureus KPR* (4S3M)- aquamarine
(F) *Pseudomonas aeruginosa* KPR2 (8IWG) – deep blue, *Methylococcus capsulatus* KPR (3I83) – light grey

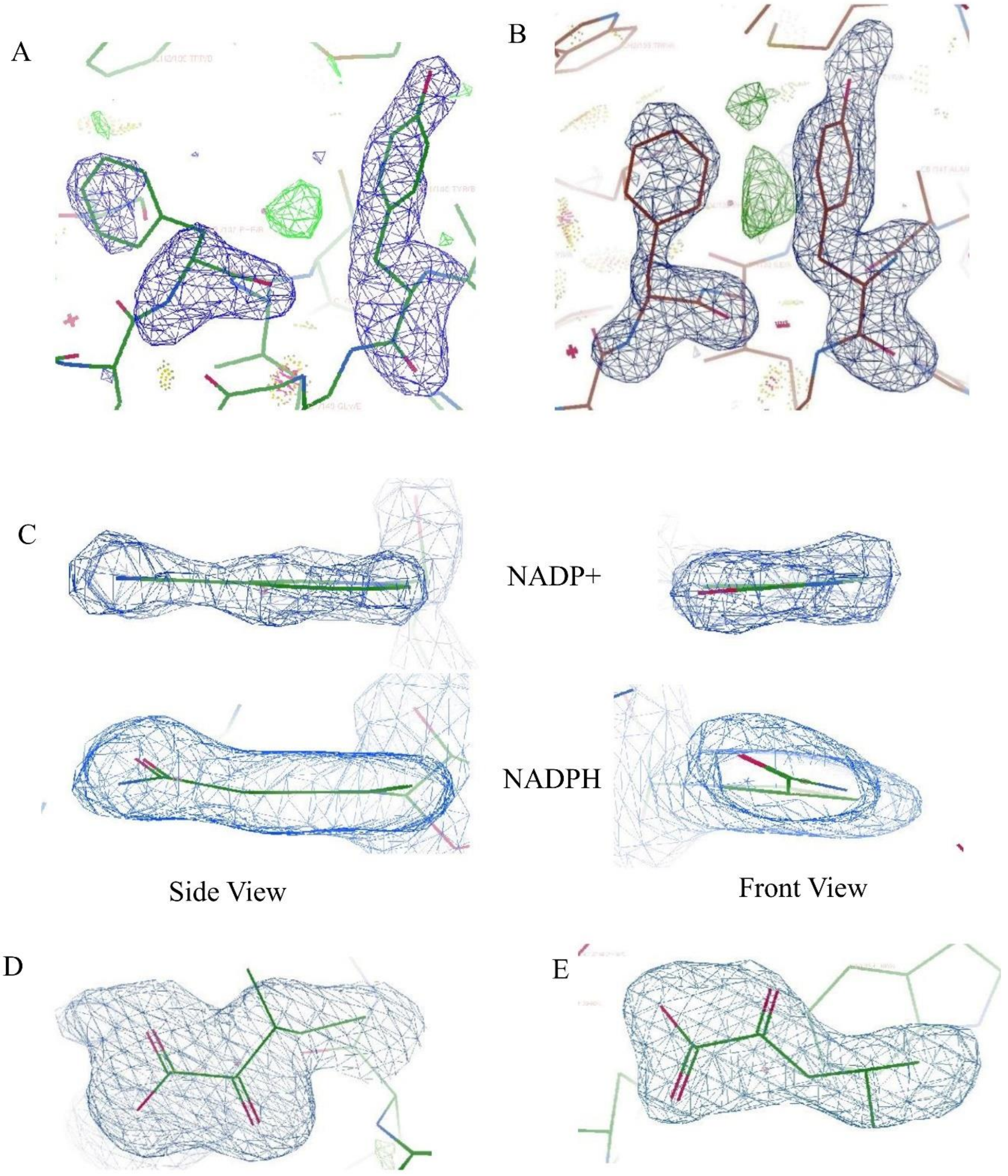


**Supplementary Figure 6:** Electron density from the KPR structures. (A,B) Respective orientation of F 132 and Y148 in closed and open molecular gate shows electron density from the final 2*F*o - *F*c map contoured at 1*ό*. (C) . Front view and side view of Polder OMIT maps captures puckered conformation of Nicotiamide group of NADPH and as compared to NADP+. (D) Polder OMIT maps of Keto-isoleucine in structure 8IX9 with 3σ contour level, (E) Polder OMIT maps of Keto-isocaproate in structure 8IXM with 3σ contour level. Ligand and residue atoms are colored according to convention and maps are displayed as blue mesh

**Supplementary Table 1: Data Collection and Refinement Statistics of the structures** (values for the outer shell given in the parenthesis).

| **PDB ID** | **8IWG** | **8IWQ** | **8IX9** | **8IXH** | **8IXM** |
|---|---|---|---|---|---|
| **Diffraction Source** | Bruker X8 Proteum | Bruker X8 Proteum | Bruker X8 Proteum | Synchrotron-RRCAT INDUS-2 BEAMLINE PX-BL21 | Bruker X8 Proteum |
| **Wavelength (Å)** | 1.54 | 1.54 | 1.54 | 0.979 | 1.54 |
| **Temperature (K)** | 100 | 100 | 100 | 100 | 100 |
| **Space group** | P 1 21 1 | P 1 21 1 | C 1 2 1 | C 1 2 1 | P 1 21 1 |
| **Unit Cell Dimensions** | | | | | |
| a, b, c (Å) | 67.1, 47.69, 96.26 | 66.85, 47.5, 96.27 | 192.23, 48.15, 128.31 | 192.21, 47.92, 129.23 | 66.59, 46.77,95.78 |
| α, β, γ (°) | 90, 99.31, 90.00 | 90.00, 99.46, 90.00 | 90.00, 129.70, 90.00 | 90.00, 129.68, 90.00 | 90.00, 98.11, 90.00 |
| Resolution range (Å) | 28.7-2.15 (2.23-2.15) | 26.86-2.19 (2.27-2.19) | 25.11-2.20 (2.28-2.20) | 40.84-1.95 (2.02-1.95) | 23.42-1.96 (2.03-1.96) |
| Total no of reflections | 345563 (32929) | 161965 (14460) | 376798 (34411) | 252569 (25373) | 386894 (36752) |
| No. of Unique reflections | 33097 (3276) | 30908 (2947) | 45956 (4378) | 66387 (6616) | 42331 (4206) |
| CC 1/2 | 0.997 (0.778) | 0.995 (0.703) | 0.995 (0.870) | 0.998 (0.632) | 0.998 (0.547) |
| Completeness (%) | 99.9 (95.2) | 98.6 (94.2) | 98.9 (93.6) | 99.9 (99.9) | 100 (100) |
| Redundancy | 10.4 (10.1) | 5.2 (4.9) | 8.2 (7.9) | 3.8 (3.8) | 9.1 (8.7) |
| I/σI | 8.6 (1.7) | 9.5 (2.3) | 11.3 (3.4) | 10.7 (1.6) | 12.0 (2.1) |
| **Refinement** | | | | | |
| R free (%) | 19.5 | 19.5 | 17.6 | 19.7 | 19.2 |
| R work (%) | 22.8 | 24.2 | 21 | 22 | 23.7 |
| Wilson Bfactor (Å$^2$) | 28.7 | 21 | 14 | 29.3 | 15.6 |
| Average Bfactor (Å$^2$) | 38 | 33 | 19 | 35 | 30 |
| **R.M.S Deviation** | | | | | |
| Bonds (Å) | 0.004 | 0.002 | 0.004 | 0.004 | 0.003 |
| Angles (°) | 0.682 | 0.553 | 0.676 | 0.776 | 0.616 |
| **Ramachandran Plot** | | | | | |
| Favoured (%) | 97.41 | 97.89 | 97.20 | 96.85 | 97.24 |
| Allowed (%) | 2.43 | 1.18 | 2.80 | 3.04 | 2.76 |
| Outlier (%) | 0.16 | 0.93 | 0.00 | 0.11 | 0.00 |
